# Principles of assembly and regulation of condensates of Polycomb repressive complex 1 through phase separation

**DOI:** 10.1101/2022.12.26.521954

**Authors:** Kyle Brown, Pin Yu Chew, Steven Ingersoll, Jorge R Espinosa, Anne Aguirre, Tatiana Kutateladze, Rosana Collepardo Guevara, Xiaojun Ren

## Abstract

PRC1 (Polycomb repressive complex 1) plays a significant role in cellular differentiation and development by repressing lineage-inappropriate genes. PRC1 proteins phase separate to form Polycomb condensates (bodies) that are multi-component hubs for silencing Polycomb target genes; however, the molecular principles that underpin the condensate assembly and biophysical properties remain unknown. Here, by using biochemical reconstitution, cellular imaging, and multiscale molecular simulations, we show that PRC1 condensates are assembled via a scaffold-client liquid–liquid phase separation (LLPS) model by which Chromobox 2 (CBX2) is the scaffold and other subunits of the CBX2-PRC1 complex act as clients. The clients induce a reentrant phase transition of CBX2 condensates in a concentration-dependent manner. The composition of the multi-component, heterotypic LLPS systems directs the assembly and biophysical properties of CBX2-PRC1 condensates and selectively promotes the formation of CBX4-PRC1 condensates, but specifically dissolves condensates of CBX6-, CBX7-, and CBX8-PRC1. Additionally, the composition of CBX2-PRC1 condensates controls the enrichment of CBX4-, CBX7-, and CBX8-PRC1 into condensates but the exclusion of CBX6-PRC1 from condensates. Our results show the composition- and stoichiometry-dependent scaffold-client assembly of multi-component PRC1 condensates and supply a conceptual framework underlying the molecular basis and dynamics of Polycomb condensate assembly.

## INTRODUCTION

The genome is compartmentalized into transcriptionally active euchromatin, constitutive heterochromatin (centromeres and telomeres), and facultative heterochromatin (silenced developmental genes and inactivated X-chromosome in females) (Trojer and Reinberg, 2007; Zylicz and Heard, 2020). Membrane-less condensates, thought to form via liquid–liquid phase separation (LLPS) of nucleic acid and protein mixtures, are involved in the formation and maintenance of these distinct chromatin compartments (Feric and Misteli, 2021; Hnisz et al., 2017; Larson and Narlikar, 2018). Dysregulation of transcriptional condensates is also associated with many diseases (Ahn et al., 2021; Li et al., 2020; Shi et al., 2021). Despite their significance, there is a paucity of information on the molecular mechanisms of assembly of transcriptional condensates of multi-component systems, preventing the development of new therapeutic strategies to target proteins and nucleic acids involved in phase separation.

The repressed states of developmental genes are maintained by the formation of facultative heterochromatin, which strongly relies on Polycomb group (PcG) complexes comprised of Polycomb repressive complex (PRC) 1 and 2 (Trojer and Reinberg, 2007; Zylicz and Heard, 2020). PRC2 is a methyltransferase that tri-methylates H3K27 to generate H3K27me3, and PRC1 is a ubiquitin ligase that mono-ubiquitinates H2AK119 to produce H2AK119ub1 (Blackledge and Klose, 2021; Schuettengruber et al., 2017). PRC1 and PRC2 coordinate to establish and maintain Polycomb domains by reinforcing each other’s activity through a feedback mechanism: H3K27me3 provides a docking site for CBX-PRC1 (canonical PRC1), and H2AK119ub1 stimulates the PRC2 activity (Blackledge and Klose, 2021; Schuettengruber *et al*., 2017). Polycomb proteins are clustered into Polycomb condensates (also termed Polycomb bodies) (Bantignies et al., 2011; Buchenau et al., 1998; Cheutin and Cavalli, 2012; Gonzalez et al., 2014; Isono et al., 2013; Ren et al., 2008; Saurin et al., 1998; Wani et al., 2016), which have been proposed to be hubs for Polycomb target gene silencing (Hodgson and Brock, 2011; Pirrotta and Li, 2012; Sievers and Paro, 2013). However, the molecular mechanisms underlying Polycomb condensate assembly are far from being understood.

It has been shown that Polycomb condensates are assembled through LLPS (Kent et al., 2020; Plys et al., 2019; Seif et al., 2020; Tatavosian et al., 2019; Wu et al., 2022). The formation of CBX-PRC1 condensates is driven by a charged disordered region of CBX2 (Kent *et al*., 2020; Plys *et al*., 2019; Tatavosian *et al*., 2019). The same charged region of CBX2 is important for proper development and differentiation (Grau et al., 2011; Jaensch et al., 2021; Lau et al., 2017). Additionally, Polycomb condensates accelerate the process of locating CBX2 onto its cognate target sites (Kent *et al*., 2020). At the molecular level, these results support that Polycomb condensates function as repressive hubs. However, the assembly of Polycomb condensates through LLPS has been questioned since concentrations of CBX-PRC1 components within Polycomb condensates are below the critical concentrations needed for LLPS in some contexts (Blackledge and Klose, 2021). To resolve this disparity, it is necessary to dissect the molecular assembly of Polycomb condensates.

Canonical PRC1 is divided into five CBX-PRC1 (CBX2/4/6/7/8-PRC1) complexes (Kim and Kingston, 2022; Piunti and Shilatifard, 2021) (**Figure 1A** and **4A**). Individual CBX-PRC1 complexes are composed of one of the CBX proteins (CBX2/4/6/7/8), one of RING1A or RING1B, one of MEL18 or BMI1, and one of the PHC proteins (PHC1/2/3). The ring finger proteins (RING1A/B) and the PCGF proteins (MEL18 and BMI1) appear as condensates (Huseyin and Klose, 2021; Kent *et al*., 2020), but how they are assembled into condensates is still unclear. The PHC proteins can polymerize (Kim et al., 2002), and their polymerization activity affects the assembly of PRC1 condensates (Isono *et al*., 2013). Polyhomeotic (Ph), the mammalian PHC counterpart in *Drosophila*, can phase separate independently of its polymerization activity (Seif *et al*., 2020). As such, the role of PHC proteins in condensate assembly is not fully understood. All CBX proteins have a conserved N-terminal Chromodomain and a conserved C-terminal Chromobox (Kim and Kingston, 2022; Piunti and Shilatifard, 2021). They share redundant, but distinct, functions during development and differentiation in a context-dependent manner (Kim and Kingston, 2022; Piunti and Shilatifard, 2021). The CBX2/4/7/8 proteins appear as condensates, while CBX6 shows uniform distribution in mouse embryonic stem (mES) cells (Ren *et al*., 2008). The molecular mechanisms underlying the condensate-forming specificity of CBX proteins are still elusive.

**Figure 1.**
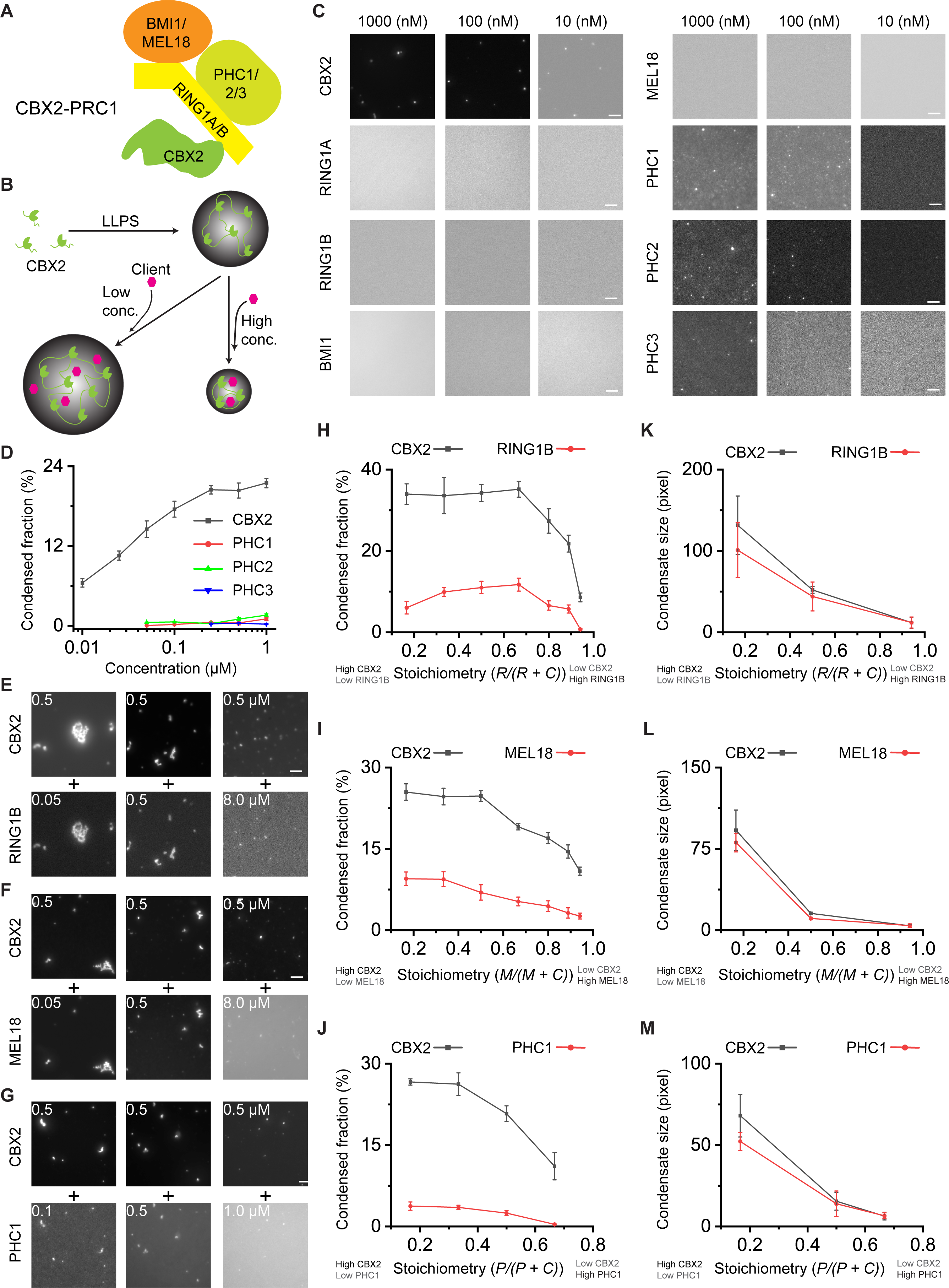
CBX2-PRC1 condensates are assembled through a scaffold-client mechanism regulated by stoichiometry. (**A**) Schematic representation of CBX2-PRC1. CBX2-PRC1 is comprised of CBX2, one RING1A or RING1B (RING1A/B), one PHC1, PHC2, or PHC3 (PHC1/2/3), and one BMI1 or MEL18. (**B**) A hypothetical model describing how CBX2-PRC1 is assembled into condensates through liquid-liquid phase separation (LLPS). Phase separated CBX2 (Chromobox 2) condensates recruit CBX2-PRC1 subunit clients that modulate the properties of condensates in a concentration-dependent manner. Clients at a relative low concentration promote the formation of CBX2 condensates, while clients at a relative high concentration dissolve CBX2 condensates. Magenta hexagon represents CBX2-PRC1 subunit clients. (**C**) Representative epi-fluorescence images of condensates of individual CBX2-PRC1 components. The CBX2-PRC1 proteins were expressed and purified from *E. Coli*. Condensates are formed in the presence of 100 mM KCl and 1.0 mM MgCl_2_ unless otherwise specified. The CBX2-PRC1 proteins were fused with fluorescent proteins. Scale bar, 5.0 μm. (**D**) Condensed fraction of CBX2 and PHC1/2/3 quantified from Figure 1C. RING1A/B and BMI1/MEL18 do not form condensates under the conditions tested. Data shown is from at least ten frames. Error bars are SD. (**E-G**) Representative epi-fluorescence images of the scaffold CBX2 and the CBX2-PRC1 subunit clients (RING1B, MEL18, and PHC1). CBX2, at a fixed concentration of 0.5 µM, was mixed with a serial of concentrations of RING1B, MEL18, and PHC1, respectively. For each panel, the top are images of CBX2 condensates while the bottom are images of the CBX2-PRC1 subunits. Scale bar, 5.0 μm. (**H-M**) Condensed fraction (**H-J**) and condensate size (**K-M**) of the scaffold CBX2 and the CBX2-PRC1 subunit clients quantified from Figure 1E-G. The black line shows the condensed fraction and condensate size of CBX2 in the presence of different concentrations of the CBX2-PRC1 subunits. The red line shows the condensed fraction and condensate size of the CBX2-PRC1 subunits in the presence of 0.5 µM CBX2. Data shown is from at least ten frames. Error bars stand for SD.

The expression level of CBX-PRC1 proteins is developmentally and pathologically regulated, forming distinct multi-protein complexes in specific developmental stages and in different diseases (Kim and Kingston, 2022; Piunti and Shilatifard, 2021), suggesting that their composition and concentration contribute to physiological and pathological conditions. Recently, it has been shown that the composition and concentration direct the assembly and biophysical properties of multi-component phase separation systems (Banani et al., 2016; Espinosa et al., 2020; Ferrolino et al., 2018; Lyon et al., 2021; Mittag and Pappu, 2022; Riback et al., 2020; Shrinivas and Brenner, 2021). Yet, whether and how the composition and concentration of CBX-PRC1 proteins contribute to the Polycomb condensate assembly is still unknown.

Here, by using a systematic approach, we obtain a holistic view of the determinants of CBX-PRC1 condensate formation and gain insights into how these components work together to assemble into CBX-PRC1 condensates. Our results show that not all CBX2-PRC1 components affect condensate stability and properties equally, but rather CBX-PRC1 condensates are assembled via a scaffold–client model, in which CBX2 is the scaffold and the other CBX2-PRC1 components are clients. Within this model, the scaffolds are the biomolecules, or groups of biomolecules, that are indispensable for condensate formation, while clients are the biomolecules recruited to the multi-component condensate via scaffold–client interactions. The composition and stoichiometry of these heterotypic LLPS systems regulate the partitioning and liquid-like behavior of CBX2-PRC1 condensates. We also show that composition-dependent heterotypic LLPS drives the specific exclusion of CBX6 from CBX2-PRC1 condensates but promotes the selective enrichment of CBX4, CBX7, and CBX8 into CBX2-PRC1 condensates. As such, we supply a biochemical and biophysical framework that underpins the assembly and properties of Polycomb condensates, which paves the avenue for further dissecting the molecular processes of Polycomb condensates in the formation and maintenance of facultative heterochromatin.

## RESULTS

### CBX2-PRC1 condensates are assembled through a scaffold-client mechanism regulated by stoichiometry

Previous biochemical studies have shown that CBX2-PRC1, one of the CBX-PRC1 complexes, is composed of CBX2, one of RING1A or RING1B, one of MEL18 or BMI1, and one of PHC1, PHC2, or PHC3 (Blackledge and Klose, 2021) (**Figure 1A**), but the composition of multi-component Polycomb condensates remains undetermined. Prior studies have shown that CBX2 drives the formation of CBX2-PRC1 condensates (Kent *et al*., 2020; Plys *et al*., 2019; Tatavosian *et al*., 2019); however, how other components are assembled into CBX2 condensates and whether they regulate the assembly remain unknown. To solve this conundrum, investigating the role of the various CBX2-PRC1 components is important. From the molecular point of view, phase separation leading to the formation of multi-component condensates is contributed by a multitude of cooperative and competitive interactions among the components and the solvent. The stoichiometry of the various components and the solution conditions (e.g., temperature, pH, type and concentration of ions) can alter the interplay of intermolecular interactions and the balance between enthalpy and entropy. hence transforming both the saturation concentrations needed to trigger phase separation and the stability of the condensates (Banani *et al*., 2016; Espinosa *et al*., 2020; Ferrolino *et al*., 2018; Lyon *et al*., 2021; Mittag and Pappu, 2022; Riback *et al*., 2020; Shrinivas and Brenner, 2021). Among the various components, only a subset of them, known as the scaffolds (Banani *et al*., 2016; Espinosa *et al*., 2020; Ferrolino *et al*., 2018; Lyon *et al*., 2021; Mittag and Pappu, 2022; Riback *et al*., 2020; Shrinivas and Brenner, 2021), seem to be indispensable to trigger condensate formation. This is because scaffold–scaffold interactions provide sufficient enthalpic gain to compensate for the entropic cost of forming the condensates. Nonetheless, other molecules recruited to the multi-component condensates, known as clients, can alter the stability of multi-component condensates in non-trivial manners: condensate stability may be decreased through competition with the scaffolds or increased by bridging scaffolds (Banani *et al*., 2016; Espinosa *et al*., 2020; Ferrolino *et al*., 2018; Lyon *et al*., 2021; Mittag and Pappu, 2022; Riback *et al*., 2020; Shrinivas and Brenner, 2021). Since CBX2-PRC1 condensates are a multi-component system, we hypothesized that CBX2-PRC1 condensates are assembled through the scaffold-client model where CBX2 is a scaffold while the other CBX2-PRC1 subunits are clients that regulate condensate assembly (**Figure 1B**).

To test this hypothesis, we first compared the ability of the separate CBX2-PRC1 subunits (i.e., CBX2, RING1A, RING1B, MEL18, BMI1, PHC1, PHC2, and PHC3) to form condensates by *in vitro* biochemical reconstitution. We expressed and purified the individual CBX2-PRC1 subunits from *E. coli* (**Figure S1A**). CBX2-PRC1 components were fused with fluorescent proteins (mCherry, YFP, or Cerulean), which allowed us to image condensates by fluorescence microscopy. We performed the condensate formation assay by using the buffer of 100 mM KCl and 1.0 mM MgCl_2_ without crowding agents or nucleic acids (unless otherwise specified, the same buffer was used throughout the study) at room temperature. In the presence of Mg^2+^, CBX2 phase separated on its own, forming single-component condensates (herein, we refer to the number of components as the number of distinct proteins in the system), at concentrations as low as 10 nM (**Figure 1C**). Previously, in buffers absent of Mg^2+^ but containing crowding agents or nucleic acids, CBX2 concentrations in the micromolar range were required to form single-component CBX2 condensates (Kent *et al*., 2020; Tatavosian *et al*., 2019). Thus, our results reveal that Mg^2+^ significantly promotes CBX2 phase separation.

At low CBX2 concentrations, small and circular condensates formed (**Figure 1C**), whereas at high CBX2 concentrations, large and irregular condensates developed (**Figure S1B**). In isolation, neither RING1A, RING1B, BMI1 nor MEL18 formed condensates under the conditions tested (i.e., up to 8 µM protein concentration). PHC1, PHC2, and PHC3 were able to form condensates but required protein concentrations one order of magnitude higher thasn required for CBX2 (i.e., equal to, or greater than, 100 nM) (**Figure 1C**). Besides the differences in protein concentrations needed to trigger phase separation, we noted that the condensed fraction and condensate size of single-component CBX2 condensates were much larger than those of the single-component PHC counterparts (**Figure 1D** and **S1C**). These results indicate that CBX2 has a greater ability to phase separate than all the other CBX2-PRC1 subunits, including the PHC family proteins, at the buffer conditions and temperature tested. Since PHC proteins can polymerize (Kim *et al*., 2002), we investigated whether their polymerization ability influences their phase behaviour. We generated PHC1^L307R^, which is defective in polymerization. PHC1^L307R^ had a similar ability to form condensates as wild-type PHC1 (**Figure S1D**). As such, we generate the first full picture of the ability of individual CBX2-PRC1 subunits to form single-component protein condensates. That is, among the various CBX2-PRC1 subunits, only CBX2 and the PHC family formed single-component condensates under the conditions assayed; however, CBX2 showed a much greater ability to phase separate than the PHC family of proteins. Therefore, we hypothesized that during the formation of multi-component Polycomb condensates, CBX2 may act as the scaffold that drives phase separation and recruits the other CBX2-PRC1 subunits as clients.

To investigate this further, we tested the phase separation capacity of CBX2 in combination with one of the other CBX2-PRC1 subunits (i.e., either RING1B, MEL18, or PHC1). To do this, we performed a new two-component condensate formation assay, where we fixed the concentration of CBX2 and varied the concentrations of one the other CBX2-PRC1 subunits. Epi-fluorescence imaging showed that while RING1B and MEL18 do not form condensates on their own, they are recruited into CBX2 condensates (**Figure 1E-F** and **S1E**). Although PHC1 could form condensates on its own, the condensed fraction of PHC1 inside two-component CBX2/PHC1 condensates was much larger than that observed in single-component PHC1 condensates (**Figure 1G** and **Figure S1F**). These results reveal that CBX2 promotes the enrichment of all CBX2-PRC1 subunits within two-component condensates, providing further evidence that CBX2 acts as the scaffold, and the other CBX2-PRC1 subunits as clients.

The formation and dissolution of condensates are fundamentally important in the regulation of condensate activities. Earlier studies have shown that varying component concentrations can mediate a reentrant liquid condensate phase of multi-component systems (Banerjee et al., 2017; Burke et al., 2015; Choi et al., 2019; Ditlev et al., 2018; Henninger et al., 2021; Milin and Deniz, 2018). A reentrant phase transition occurs when the monotonic variation of a single thermodynamic parameter (e.g., client concentration, RNA concentration, salt concentration) induces a change in the system from one thermodynamic state to a macroscopically similar state via two-phase transitions (Krainer et al., 2021). For instance, in the well-known RNA-driven reentrant phase behavior of RNA binding proteins such as FUS, increasing RNA concentration initially enhances the stability of RNA/protein condensates, but when the concentration of RNA surpasses a threshold, the stability of the condensates begins to decrease until, eventually, dissolution is triggered (Banerjee *et al*., 2017). Similarly, in the two-component CBX2/RING1B systems we studied, our quantitative results show that when RING1B is present in concentrations smaller than that of CBX2, RING1B promotes the formation of CBX2 condensates; however, when the RING1B concentration exceeds that of CBX2, RING1B dissolves the two-component condensates (**Figure 1H-M** and **S1F**). In two-component systems of CBX2 and either MEL18 or PHC1, MEL18 and PHC1 promoted the formation of CBX2 condensates when present at a concentration lower than the equimolar ratio; however, their ability to promote condensates was modest when compared to that of RING1B (**Figure 1H-M** and **S1F**). When the concentration of MEL18 and PHC1 was higher than the equimolar mixture, they also dissolved CBX2 condensates (**Figure 1H-M**).

Our results show that CBX2 phase separates into condensates and suggest that CBX2 acts as a scaffold that recruits CBX2-PRC1 subunits as clients. Our results also postulate that CBX2-PRC1 subunit clients induce a previously undescribed, concentration-dependent reentrant phase transition of CBX2 condensates—where a relatively low concentration of clients promotes the formation of CBX2 condensates, while a relatively high concentration of clients dissolves CBX2 condensates (**Figure 1B**).

### Cellular imaging supports a scaffold-client model and a client-inclusive regulatory mechanism

The scaffold-client model for the regulation of multi-component CBX2-PRC1 condensates would imply that CBX2 proteins should be enriched within such condensates in cells. Earlier studies have shown that overexpressed CBX2 in cells promotes the formation of condensates (Kent *et al*., 2020; Plys *et al*., 2019; Ren *et al*., 2008; Tatavosian *et al*., 2019); however, it still is unknown whether endogenous CBX2 forms condensates at physiological concentrations. To this end, we engineered a HaloTag (HT) at the C-terminus of endogenous *Cbx2* by using CRISPR/Cas9 (Ran et al., 2013), generating homozygous *Cbx2^HT/HT^* and heterozygous *Cbx2^WT/HT^* knock-in mouse embryonic stem (mES) cell lines (**Figure 2A-C**). Following labelling with the HaloTag TMR ligand, epi-fluorescence imaging showed that HT-CBX2 in homozygous *Cbx2^HT/HT^* and heterozygous *Cbx2^WT/HT^* exhibit granular distributions in living cells, with approximately 100 condensates per cell (**Figure 2D**). The intensity ratio of the dense to dilute phase was about 1.5. The condensed fraction was approximately 20%. Co-immunofluorescence showed that PHC1 and RING1B were concentrated inside the CBX2 condensates in *Cbx2^HT/HT^* knock-in mES cells (**Figure 2E** and **S2A**). We also randomly integrated *FLAG-HaloTag-CBX2* (*F-HT-CBX2*) into the genome of HeLa cells. Co-immunofluorescence showed that RING1B, BMI1, PHC1, and PHC1, as well as H3K27me3 are all enriched within CBX2 condensates (**Figure S2B-C**). As such, our results show that in living cells, CBX2 forms condensates enriched in the CBX2-PRC1 client subunits.

**Figure 2.**
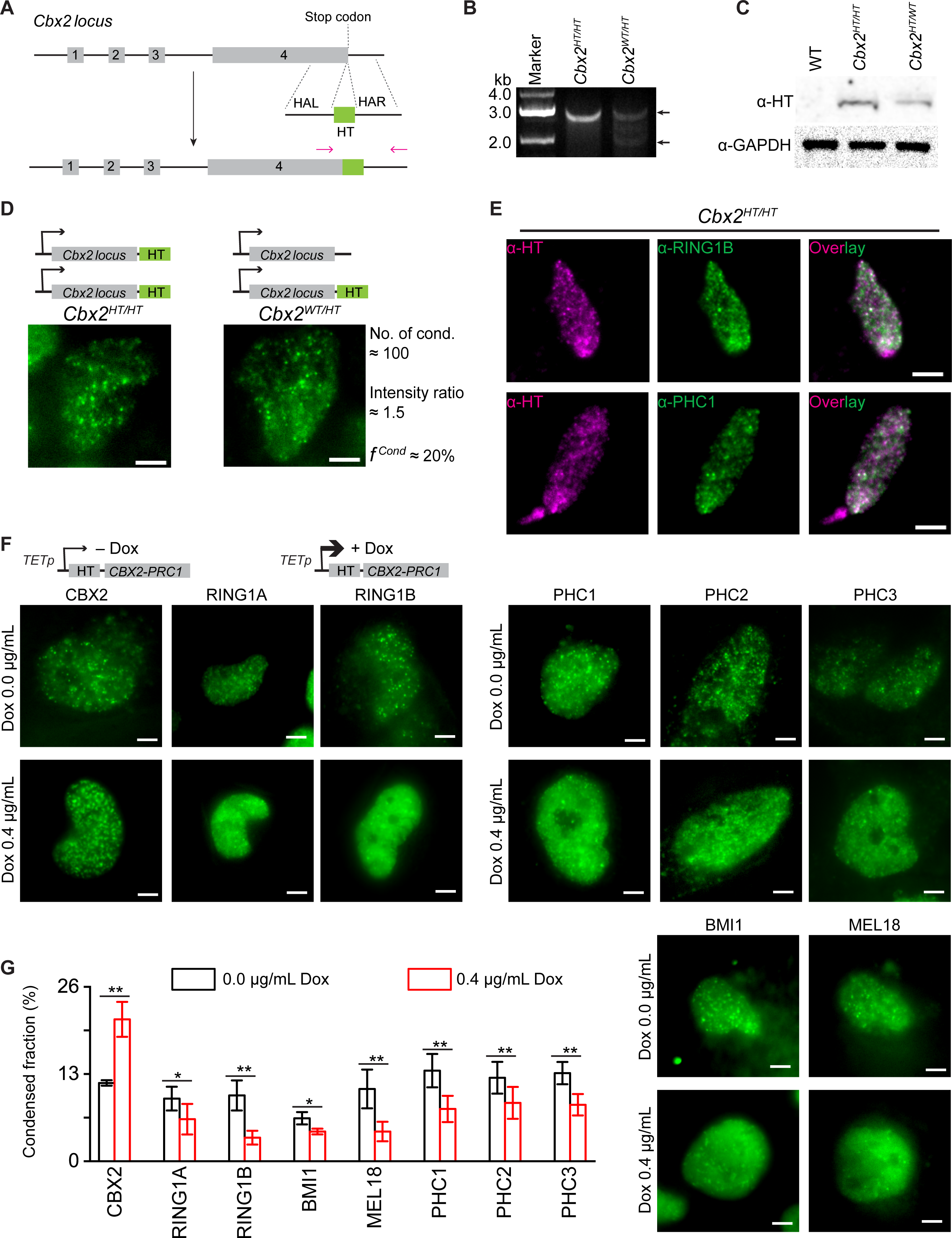
Cellular imaging supports a scaffold-client model and a client-inclusive regulatory mechanism. (**A**) Schematic representation for CRISPR/Cas9-mediated homologous recombination to insert HaloTag to the C-terminus of *Cbx2* in mouse embryonic stem (mES) cells. The red arrowheads indicate the primers used to verify the insertion. HAL, homology left arm; HAR, homology right arm; and HT, HaloTag. (**B**) Agarose gel analysis of PCR amplicons from homozygous HaloTag (*Cbx2^HT/HT^*) and heterozygous HaloTag (*Cbx2^WT/HT^*) knockin mES cell lines. The arrowheads show the correct size of PCR amplicons. The PCR primers used are shown in Figure 2A. (**C**) Western blots for HaloTag comparing wild-type (WT), homozygous CBX2-HT, and heterozygous CBX2-HT mES cell lines. Glyceraldehyde-3-phosphate dehydrogenase (GAPDH) was the loading control. (**D**) Live-cell imaging subnuclear distribution of CBX2-HaloTag (CBX2-HT) in homozygous *Cbx2^HT/HT^* (left panel) and heterozygous *Cbx2^HT/WT^* (right panel) mES cell lines. CBX2-HT mES cells are labelled with HaloTag® TMR ligand. The right shows the number of condensates, the intensity ratio of condensates to non-condensed regions, and the condensed fraction, which are similar for both cell lines. Scale bar, 5.0 µm. (**E**) Co-immunostaining analysis of CBX2-HT as well as endogenous RING1B and PHC1 in *Cbx2^HT/HT^* cell line. The right panel shows image overlay. CBX2-HT was stained by anti-HaloTag antibody. RING1B and PHC1 were stained by anti-RING1B and anti-PHC1 antibodies, respectively. Scale bar, 5.0 µm. (**F**) Live-cell imaging subnuclear localization of the components of CBX2-PRC1. The CBX2-PRC1 component fused with HaloTag were stably integrated into the genome of HeLa cells. The expression level is controlled by a tetracycline-response element (top panel) and induced by doxycycline (Dox). The basal expression was without adding Dox. The expression was induced by 0.4 µg/mL Dox. The thinness of arrowhead corresponds to the level of expression. The HaloTag fusion proteins were labelled with HaloTag® TMR ligand. Scale bar, 5.0 µm. (**G**) Condensed fraction of the components of CBX2-PRC1 quantified from Figure 2F. The results shown are representative of data from at least ten cells (*, P < 0.05; **, P < 0.01). Error bars are SD.

Our *in vitro* biochemical reconstitution suggests that the relative concentration of CBX2 proteins versus that of the other CBX2-PRC1 subunits acting as clients affects the stability of the two-component CBX2-client condensates (**Figure 1H-M**). Increasing the CBX2 to client ratio should increase the condensed fraction. On the other hand, decreasing the CBX2 to client ratio should decrease the condensed fraction. To test this in living cells, we generated HeLa cell lines that stably express one of the various CBX2-PRC1 proteins fused with HaloTag, respectively.

Their expression level was controlled by a tetracycline-response promoter and induced by doxycycline (Dox) (**Figure 2F** and **S2D**). Live-cell epi-fluorescence imaging showed that all CBX2-PRC1 proteins show granular distributions (**Figure 2F**). The condensed fraction of the scaffold CBX2 protein and the client subunits were distinct in response to varying Dox concentrations (**Figure 2G**). The condensed fraction of CBX2 at the high protein level (0.4 µg/mL) was significantly larger than that at the low protein level (0.0 µg/mL), and CBX2 was the only protein to show this behavior. On the other hand, the condensed fraction of clients at high client protein levels were significantly smaller than that at lower protein levels. Thus, our cellular results support the postulate that formation of two-component CBX2-PRC1 condensates is driven by CBX2 proteins, which behave as the scaffolds, and regulated in a concentration-dependent manner by the rest of the CBX2-PCR1 subunits, which act as clients.

### Composition directs the partitioning and liquid-like properties of CBX2-PRC1 condensates

By studying the phase behavior of two-component systems containing CBX2 proteins and one of the other CBX2-PRC1 subunits, we have established the scaffold-client model in which CBX2 is the scaffold of condensates, which recruits individual CBX2-PRC1 subunits. We next investigated whether multiple components are simultaneously assembled into CBX2 condensates and whether they regulate the liquid-like properties of the condensates they form. Among the macroscopic liquid-like properties of condensates are their spherical shapes and short recovery times from fluorescence recovery after photobleaching (FRAP). From the molecular point of view, such liquid-like properties imply that molecules inside the condensates exhibit frequent binding and unbinding, free molecular diffusion, and exchange of in and out of the condensates. Accumulating evidence indicates that the liquid-like properties of multi-component condensates can be sensibly modulated by the composition and stoichiometry of the various components within (Banani *et al*., 2016; Ditlev *et al*., 2018; Espinosa *et al*., 2020; Riback *et al*., 2020); thus, we hypothesized that the condensate composition regulates the partitioning of the different components and the liquid-like properties of CBX2-PRC1 condensates (**Figure 3A**).

**Figure 3.**
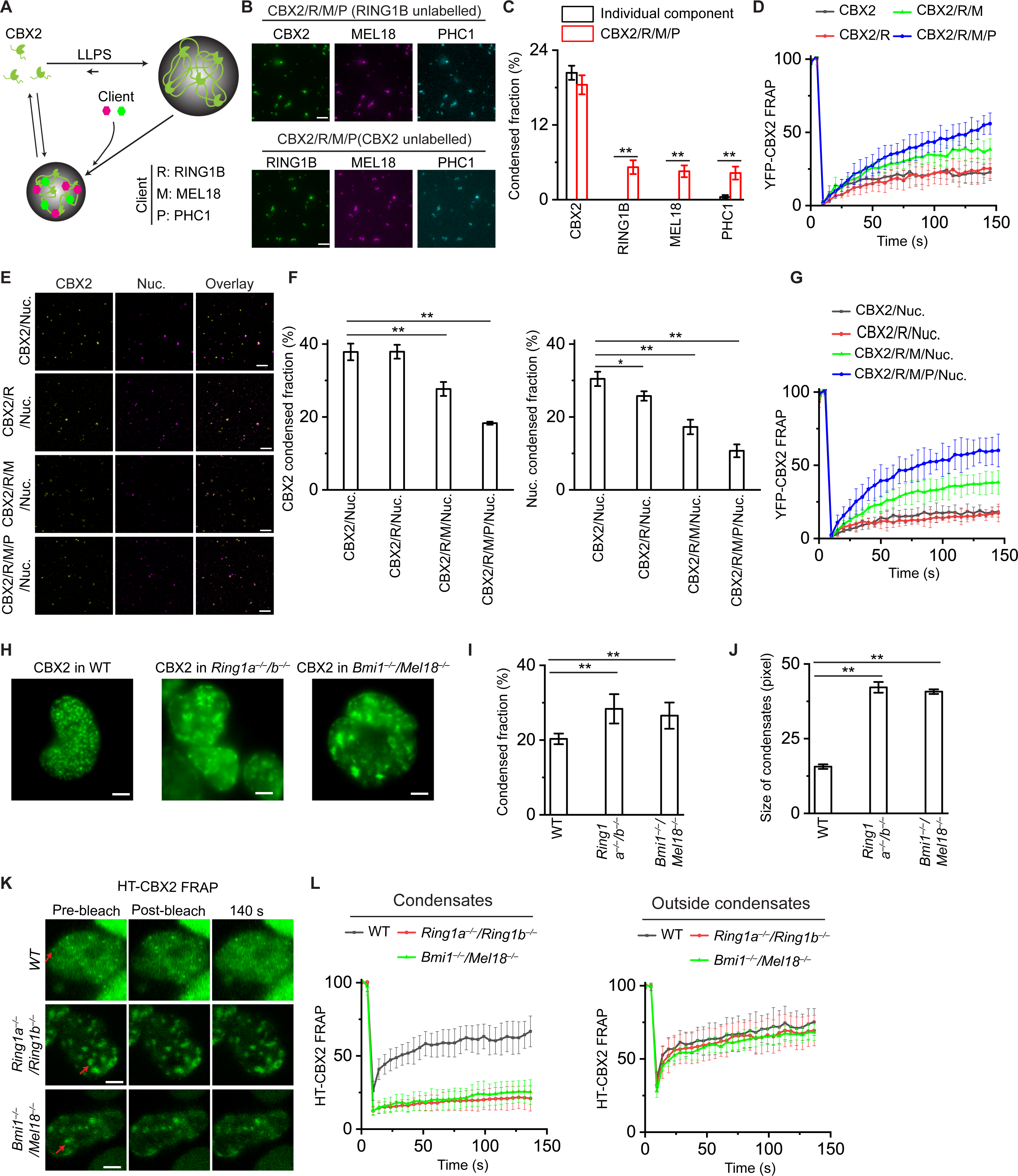
Composition directs the partitioning and liquid-like properties of CBX2-PRC1 condensates. (**A**) A hypothetical model describing how the condensate composition regulates the liquid-like properties and partitioning of CBX2-PRC1 subunit clients and nucleosomes. The inclusion of all CBX2-PRC1 components reduces partitioning of the CBX2-PRC1 subunit clients and nucleosomes into condensates but enhances the liquid-like properties of CBX2 and nucleosomes. Colored hexagons are the CBX2-PRC1 subunit clients (magenta) and nucleosomes (green). (**B**) Representative epi-fluorescence images of CBX2-PRC1 subunits in the four-component (CBX2, RING1B (R), MEL18 (M), and PHC1 (P)) system. Top panel: RING1B was un-labelled, and others are fused with different fluorescent proteins. Bottom panel: CBX2 was un-labelled, and others are fused with different fluorescent proteins. The concentration of individual components is 0.5 µM. Scale bar, 5.0 μm. (**C**) Condensed fraction of the components of CBX2-PRC1 in the four-component (CBX2/R/M/P) system quantified from Figure 3B compared with individual components in the single-component system from Figure 1D. Data are from at least ten frames (**, P < 0.01). Error bars are SD. (**D**) FRAP curves of CBX2 in the single-component (CBX2, black line), two-component (CBX2/R, red line), three-component (CBX2/R/M, green line), and four-component (CBX2/R/M/P, blue line) systems. Data shown is from at least ten condensates. Error bars stand for SD. (**E**) Example confocal fluorescence images of CBX2 and nucleosomes in the two-component (CBX2/Nuc. (nucleosomes)), three-component (CBX2/R/Nuc.), four-component (CBX2/R/M/Nuc.), and five-component (CBX2/R/M/P/Nuc. systems. CBX2 was labelled with YFP and nucleosomes with Cy3. Other CBX2-PRC1 subunits are un-labelled. The concentration of individual components is 0.5 µM and the concentration of nucleosomes is 0.3 µM. Scale bar, 5.0 μm. (**F**) Condensed fraction of CBX2 and nucleosomes in the two-component, three-component, four-component, and five-component systems quantified from Figure 3E. The results shown are representative of data from at least five frames (*, P < 0.05; **, P < 0.01). Error bars are SD. (**G**) FRAP curves of YFP-CBX2 and Cy3-nucleosomes in the two-component, three-component, four-component, and five-component systems. Data shown is from at least ten condensates. Error bars are SD. (**H**) Representative live-cell epi-fluorescence images of HT-CBX2 in wild-type (WT), *Ring1a^−/−^/b^−/−^*, *Bmi1^−/−^/Mel18^−/−^* mES cell lines. HT-CBX2x induced by 0.4 µg/mL Dox was labelled with HaloTag® TMR ligand. Scale bar, 5.0 µm. (**I-J**) Condensed fraction (**I**) and size (**J**) of HT-CBX2 condensates in wild-type (WT), *Ring1a^−/−^/b^−/−^*, *Bmi1^−/−^/Mel18^−/−^* mES cell lines quantified from Figure 3H. The results shown are representative of data from at least ten cells (**, P < 0.01). Error bars are SD. (**K**) Example confocal images of FRAP of HT-CBX2 in wild-type (WT), *Ring1a^−/−^/b^−/−^*, *Bmi1^−/−^/Mel18^−/−^* mES cell lines. HT-CBX2 was labelled with HaloTag® TMR ligand. Red arrowhead shows condensates to be bleached. (**L**) FRAP curves of HT-CBX2 within and outside condensates in wild-type (WT), *Ring1a^−/−^/b^−/−^*, *Bmi1^−/−^/Mel18^−/−^* mES cell lines. Data shown is from at least five cells. Error bars are SD.

To investigate whether multiple components are recruited into CBX2 condensates, we biochemically reconstituted the four-component system by mixing YFP-CBX2 with mCherry-MEL18, Cerulean-PHC1, and unlabeled RING1B, or by mixing unlabeled CBX2 with mCherry-MEL18, Cerulean-PHC1, and YFP-RING1B. The condensed fraction of CBX2 in the four-component system was similar to that in the single-component system (**Figure 3B-C**). RING1B and MEL18 were significantly enriched within CBX2 condensates. The condensed fraction of PHC1 in the four-component system was 10-fold more than that on the PHC1 single-component condensates, and approximately 2-fold more than that in the two-component (CBX2 and PHC1) system (**Figure 3C** and **S1F**), suggesting that RING1B and MEL18 facilitate partitioning of PHC1 into the multi-component CBX2-PRC1 condensates. Since earlier studies have shown that the PHC proteins affect the assembly of condensates, we investigated whether PHC1 affects the partitioning of CBX2, RING1B, and MEL18 by reconstituting the three-component system having YFP-CBX2, Cerulean-RING1B, and mCherry-MEL18. Adding PHC1 to the three-component system significantly reduced the condensed fraction of CBX2, RING1B, and MEL18 (**Figure S3A-B**). As such, our results show that PHC1 can inhibit partitioning of RING1B and MEL18 into condensates while RING1B and MEL18 promote partitioning of PHC1 into condensates.

We next investigated whether the number of components in the CBX2 condensates regulates their liquid-like properties via FRAP (**Figure 3D** and **S3C**). CBX2 in the single-component and two-component CBX2/RING1B systems showed slow exchange with the surrounding environment. The liquid-like properties of condensates originate from weak and transient multivalent attractive interactions among the biomolecules within them. Therefore, slow exchange is indicative of relative strong CBX2–CBX2 and CBX2–RING1B interactions. Upon adding MEL18 to the two-component CBX2/RING1B system, the exchange rates of CBX2 were increased. Adding PHC1 to the three-component CBX2/RING1B/MEL18 system further increased the liquid-like behavior of CBX2. Increased exchange of CBX2 within these multi-component condensates suggests that CBX2–MEL18 and CBX2–PHC1 interactions are less energetically favorable than those sustaining the two-component CBX2/RING1B systems. As such, these results show that the condensate composition directs the liquid-like behavior of CBX2.

Earlier studies have shown that CBX2 condensates can concentrate nucleosomes (Plys *et al*., 2019; Tatavosian *et al*., 2019), but it still is unknown whether the composition of multi-component CBX2-PRC1 condensates regulates the partitioning and dynamics of nucleosomes. To this end, we reconstituted H3K27me3-mononucleosomes labeled with Cy3. We mixed Cy3-nucleosomes with YFP-CBX2 supplemented with un-labelled CBX2-PRC1 subunits. Confocal fluorescence imaging showed that nucleosomes are enriched and colocalized within the condensates (**Figure 3E** and **S3D**). Upon gradually increasing the number of components, the condensed fraction of nucleosomes was decreased with the decreased condensed fraction of CBX2 (**Figure 3F**), while CBX2 and nucleosomes became more fluid (**Figure 3G** and **S3E**). Our results show the composition of CBX2-PRC1 condensates dictates the partitioning and dynamics of nucleosomes.

Overall, our *in vitro* reconstitution shows that CBX2 condensates concentrate clients of CBX2-PRC1 subunits and nucleosomes. These clients regulate the partitioning and liquid-like behavior of components of CBX2-PRC1 condensates. As such, our results suggest a composition-dependent assembly of multi-component CBX2-PRC1 condensates.

### Cellular imaging supports that composition directs the partitioning and liquid-like properties of CBX2-PRC1 condensates

Our results suggest a model by which the composition and relative concentration of components direct the assembly and properties of CBX2-PRC1 condensates. To test whether this principle can be recapitulated in cellular conditions, we stably expressed HT-CBX2 in wild-type as well as in *Ring1a^-/-^/b^-/-^* double knockout and *Bmi1^-/-^/Mel18^-/-^* double knockout mES cells. RING1B and MEL18 are highly expressed in mES cells compared to CBX2 (Lau *et al*., 2017; Morey et al., 2012), suggesting that depletion of RING1B and MEL18 should increase the condensed fraction and condensate size of CBX2. Live-cell epi-fluorescence imaging showed that CBX2 forms small, circular condensates in wild-type mES cells but large and irregular condensates in *Ring1a^-/-^/b^-/-^* and *Bmi1^-/-^Mel18^-/-^* mES cells (**Figure 3H**), which is consistent with earlier reports (Kent *et al*., 2020; Tatavosian *et al*., 2019). The condensed fraction and condensate size of CBX2 in wild-type mES cells were significantly smaller than that in the double knockout mES cell lines (**Figure 3I-J**), indicating that CBX2-PRC1 subunits regulate the partitioning and material properties of CBX2-PRC1 condensates.

To investigate whether the composition affects the liquid-like properties of condensates, we performed FRAP of HT-CBX2 in wild-type, *Ring1a^-/-^/b^-/-^*, and *Bmi1^-/-^/Mel18^-/-^* mES cells (**Figure 3K**). CBX2 within condensates was liquid-like in wild-type mES cells (**Figure 3L**), which is consistent with earlier reports (Kent *et al*., 2020; Plys *et al*., 2019; Ren *et al*., 2008; Tatavosian *et al*., 2019). On the other hand, the recovery rate of CBX2 within condensates in *Ring1a^-/-^/b^-/-^* and *Bmi1^-/-^/Mel18^-/-^* mES cells was significantly slower than that in wild-type mES cells (**Figure 3L**). The recovery rates of CBX2 outside condensates were similar between wild-type and knockout cell lines (**Figure 3L**). As such, these cellular results support the composition-dependent control of the liquid-like properties of CBX2-PRC1 condensates.

### CBX2- and CBX4-PRC1 forms condensates *in vitro* while CBX6/7/8-PRC1s have a negligible ability to form condensates *in vitro*

We have uncovered the molecular mechanism that underpins how multi-component CBX2-PRC1 condensates are assembled and regulated. Canonical PRC1 comprises five CBX-PRC1 complexes (CBX2/4/6/7/8-PRC1s) (Blackledge and Klose, 2021) (**Figure 4A**). Earlier studies have shown that CBX4/7/8 are assembled into condensates in cells but CBX6 does not appear as condensates (Ren *et al*., 2008). The molecular mechanisms that underpin how CBX4/7/8-PRC1 condensates are assembled and why CBX6-PRC1 cannot assemble into condensates are unclear. To this end, we first investigated whether the individual CBX proteins can phase separate into condensates by biochemical reconstitution. We expressed and purified the CBX4/6/7/8 proteins fused with mCherry from *E. coli*, respectively (**Figure S4A**), and compared their condensation abilities against CBX2 by performing the condensate formation assay. Epi-fluorescence imaging showed that the CBX4/6/7/8 proteins all form condensates but with distinct abilities (**Figure S4B**). Analysis of the condensed fraction and the condensate size showed that CBX2 has the best ability to coacervate among the CBX proteins and CBX4 is better than the CBX6/7/8 proteins which have similar condensation abilities (**Figure 4B** and **S4C**). As such, our results show that all CBX proteins can form condensates *in vitro* but with distinct abilities.

**Figure 4.**
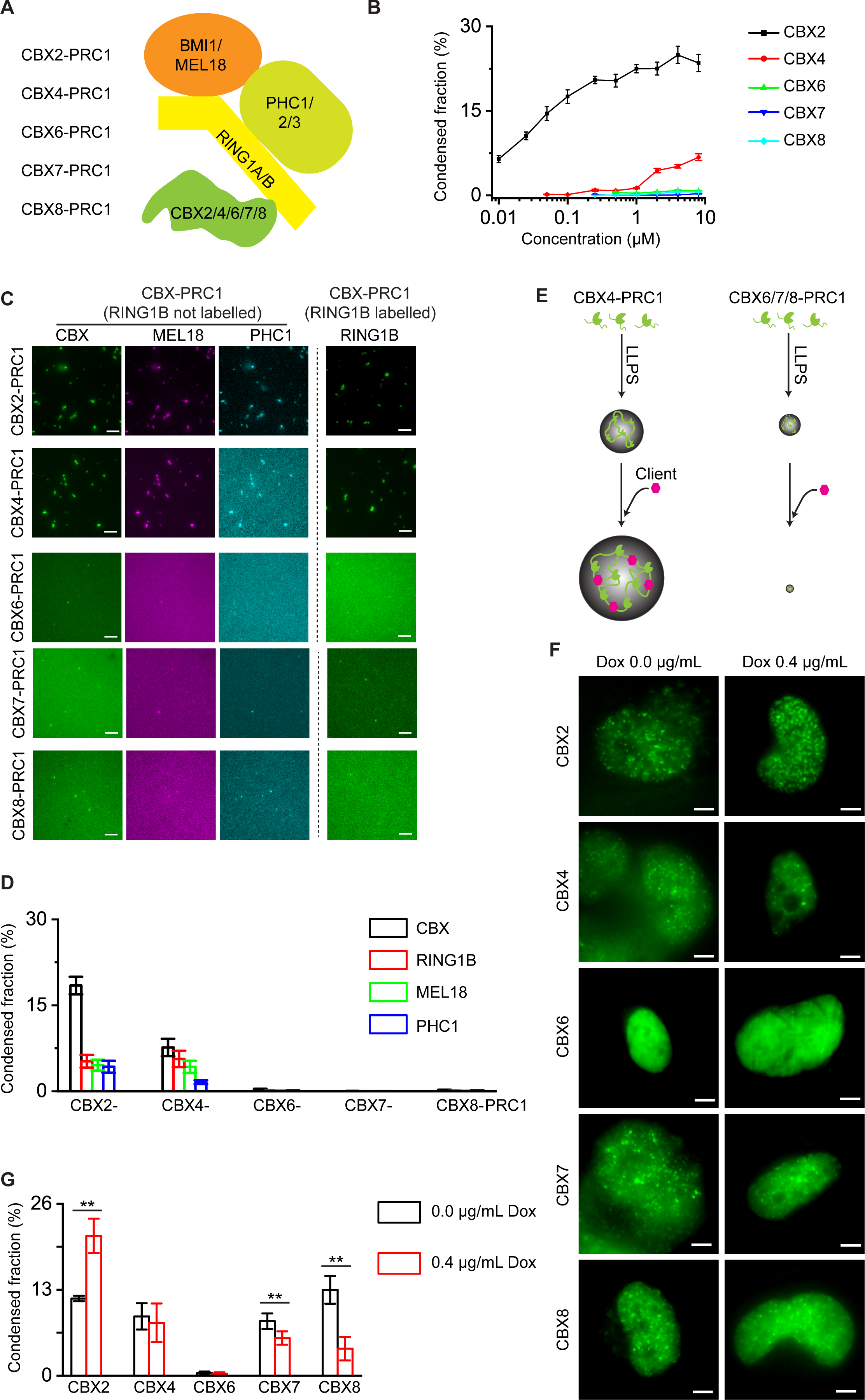
*In vitro* and *in vivo* condensation abilities of the family of CBX-PRC1 complexes. (**A**) Schematic representation of the CBX-PRC1 complexes. Canonical CBX-PRC1 is comprised of five CBX-PRC1 complexes, including CBX2-PRC1, CBX4-PRC1, CBX6-PRC1, CBX7-PRC1, and CBX8-PRC1. (**B**) Condensed fraction of the CBX proteins quantified from Figure S4B. Data shown is from at least ten frames. Error bars are SD. (**C**) Representative epi-fluorescence images of the components of CBX-PRC1 (one of CBX2/4/6/7/8, RING1B, MEL18, and PHC1). CBX2-PRC1 is replicated from Figure 3B. Panel left of the dashed line: CBX4/6/7/8 proteins were labelled with mCherry; MEL18 with YFP; and PHC1 with Cerulean. RING1B was un-labelled and not shown in the figure. Panel right of the dashed line: CBX4/6/7/8 proteins were labelled with mCherry; RING1B with Cerulean; and MEL18 with YFP. PHC1 was un-labelled. Only the RING1B images are shown. The concentration of individual components is 0.5 µM. Scale bar, 5.0 μm. (**D**) Condensed fraction of the components of CBX-PRC1 quantified from Figure 4C. Data shown is from at least ten frames. Error bars are SD. (**E**) A hypothetical model describing how individual CBX-PRC1 complexes are assembled to condensates *in vitro*. CBX4 is assembled into condensates that recruit the CBX4-PRC1 subunit clients that in turn promotes the condensation of CBX4-PRC1. The CBX6/7/8 proteins have a limited ability to form condensates. Adding CBX-PRC1 subunits dissolves CBX6/7/8 condensates. (**F**) Live-cell epi-fluorescence imaging subnuclear localization of the CBX proteins with and without Dox. The HT-CBX proteins were stably expressed in HeLa cells and labelled with HaloTag® TMR ligand. Scale bar, 5.0 µm. (**G**) Condensed fraction of the CBX proteins quantified from Figure 4F. The results shown are representative of data from at least ten cells (**, P < 0.01). Error bars are SD.

Since our results have revealed that CBX2-PRC1 condensates are assembled in a composition-dependent manner, we wondered whether composition also regulates the assembly of multi-component CBX4/6/7/8-PRC1 condensates. To this end, we biochemically reconstituted the four-component system by combining equal molar (0.5 µM) mCherry-CBX proteins with YFP-MEL18, Cerulean-PHC1, and unlabeled RING1B (RING1B not shown) (**Figure 4C, left**), or with Cerulean-RING1B, YFP-MEL18, and unlabeled PHC1 (only RING1B shown) (**Figure 4C, right**). CBX4 condensates recruited RING1B, MEL18, and PHC1. Interestingly, unlike the assembly of CBX2-PRC1 condensates, the condensed fraction of CBX4 in the four-component system (CBX4/RING1B/MEL18/PHC1) was nearly 10-fold larger than that on its own (**Figure S4D**), suggesting the composition-dependent promotion of the assembly of CBX4-PRC1 condensates. On the contrary, the condensed fraction of CBX6/7/8 in the four-component system was nearly undetectable (**Figure 4D**), suggesting a composition-dependent inhibition of the assembly of CBX6/7/8-PRC1 condensates. These biochemical reconstitution results suggest that the composition in the multi-component system promotes or inhibits the assembly of multi-component CBX-PRC1 condensates in a CBX-specific manner (**Figure 4E**).

### CBX4/7/8 proteins appear as condensates in cells but CBX6 does not

Our *in vitro* biochemical reconstitution generated a full, complex picture of the assembly of CBX-PRC1 condensates where CBX4-PRC1 is assembled into condensates while CBX6/7/8-PRC1s have nearly undetectable abilities to form condensates. Earlier live-cell results have shown that the CBX4/7/8 proteins appear as condensates and CBX6 does not appear as condensates (Ren *et al*., 2008). Thus, the biochemical reconstitution is not fully reconciled with the cellular data. To find the molecular factors that can reconcile both *in vitro* and *in vivo* data, we investigated whether the condensate assembly of the CBX proteins occurs in a concentration-dependent manner, as a hallmark of phase separation is concentration-dependence. To this end, we integrated *HT-CBX* fusions into the genome of HeLa cells, respectively. The expression level of these fusion proteins was controlled by a tetracycline-response promoter and induced by Dox (**Figure S4E**). CBX6 did not form condensates in living cells while the CBX4/7/8 proteins formed condensates (**Figure 4F**), consistent with earlier reports (Ren *et al*., 2008). Quantitative analysis of images showed that the condensed fractions of CBX4 are similar among different protein concentrations (**Figure 4G**). Larger condensates of CBX4 at the high protein concentration were seen in comparison with those at the low protein concentration (**Figure 4F**). The condensed fraction of CBX7/8 proteins at the high expression level was significantly lower than that in the low expression level (**Figure 4G**), which is similar to CBX2-PRC1 subunit clients whose condensed fractions are decreased upon increasing their concentrations. This suggests that CBX7-PRC1 and CBX8-PRC1 may act as clients.

In summary, the *in vitro* biochemical reconstitution shows that the family of CBX proteins can form condensates on their own with distinct abilities and that the composition of the multi-component system directs the assembly of CBX-PRC1 condensates *in vitro* (**Figure 4E**). The inclusion of three CBX-PRC1 subunits (RING1B/MEL18/PHC1) promotes the condensation of CBX4 but dissolves CBX6/7/8 condensates. The cellular results show that increasing concentrations of CBX7 and CBX8 decreases their condensed fraction. Thus, the *in vitro* and *in vivo* results suggest that CBX7-PRC1 and CBX8-PRC1 do not phase separate and act as clients that are recruited into condensates. CBX4-PRC1 phase separates *in vitro* and larger CBX4 condensates are seen at high protein concentration *in vivo;* however, the CBX4 condensed fractions are similar between different protein concentrations, suggesting that CBX4-PRC1 may, in part, act as a client that is recruited into condensates. CBX6-PRC1 does not form condensates both *in vitro* and *in vivo*, suggesting that CBX6-PRC1 is not recruited into condensates in cells. As such, our results suggest that CBX4//7/8-PRC1s may act as clients that are recruited into condensates and that CBX6-PRC1 is not recruited into condensates.

### CBX2-PRC1 condensates recruit CBX4/7/8-PRC1s but have a limited ability to recruit CBX6-PRC1 in a composition-dependent manner

We hypothesized that CBX2-PRC1 condensates recruit CBX4/7/8-PRC1s but cannot enrich CBX6-PRC1 (**Figure 5A**). To this end, we first asked whether CBX2 condensates can selectively recruit the family of CBX proteins. We mixed CBX2 and the other CBX proteins at an equal molar concentration (0.5 µM), respectively. CBX2 condensates were able to greatly enrich all other CBX proteins (**Figure S5A-B**), suggesting that the two-component system of CBX2 and other CBX proteins cannot confer the enrichment selectivity of CBX2 vs other CBX proteins. Since the composition of CBX2-PRC1 condensates regulates assembly, we wondered if the inclusion of other components in the system will confer enrichment and exclusion specificity of CBX2-PRC1 condensates. To this end, we investigated whether CBX2-PRC1 condensates can enrich other CBX-PRC1s by reconstituting five-component systems through mixing YFP-CBX2 with other mCherry-CBX proteins in the presence of RING1B, MEL18, and PHC1. Epi-fluorescence imaging showed that CBX6 does not appear as condensates whereas the CBX4/7/8 proteins form condensates that localize with CBX2 condensates (**Figure 5B**). The condensed fraction of CBX2 in the five-component system was approximately 6% (**Figure 5C**), which is smaller than that of CBX2 on its own (∼20%) or in the four-component system (∼20%), further supporting that composition directs the assembly of CBX-PRC1 condensates. Although CBX7-PRC1 and CBX8-PRC1 do not form condensates on their own, they were enriched within multi-component CBX2-PRC1 condensates, suggesting that CBX7-PRC1 and CBX8-PRC1 are clients of CBX2-PRC1 condensates. CBX4-PRC1 was also enriched within CBX2-PRC1 condensates. As such, our *in vitro* reconstitution shows that CBX2-PRC1 condensates selectively recruit CBX4/7/8-PRC1s but cannot enrich CBX6-PRC1 in a composition-dependent manner.

**Figure 5.**
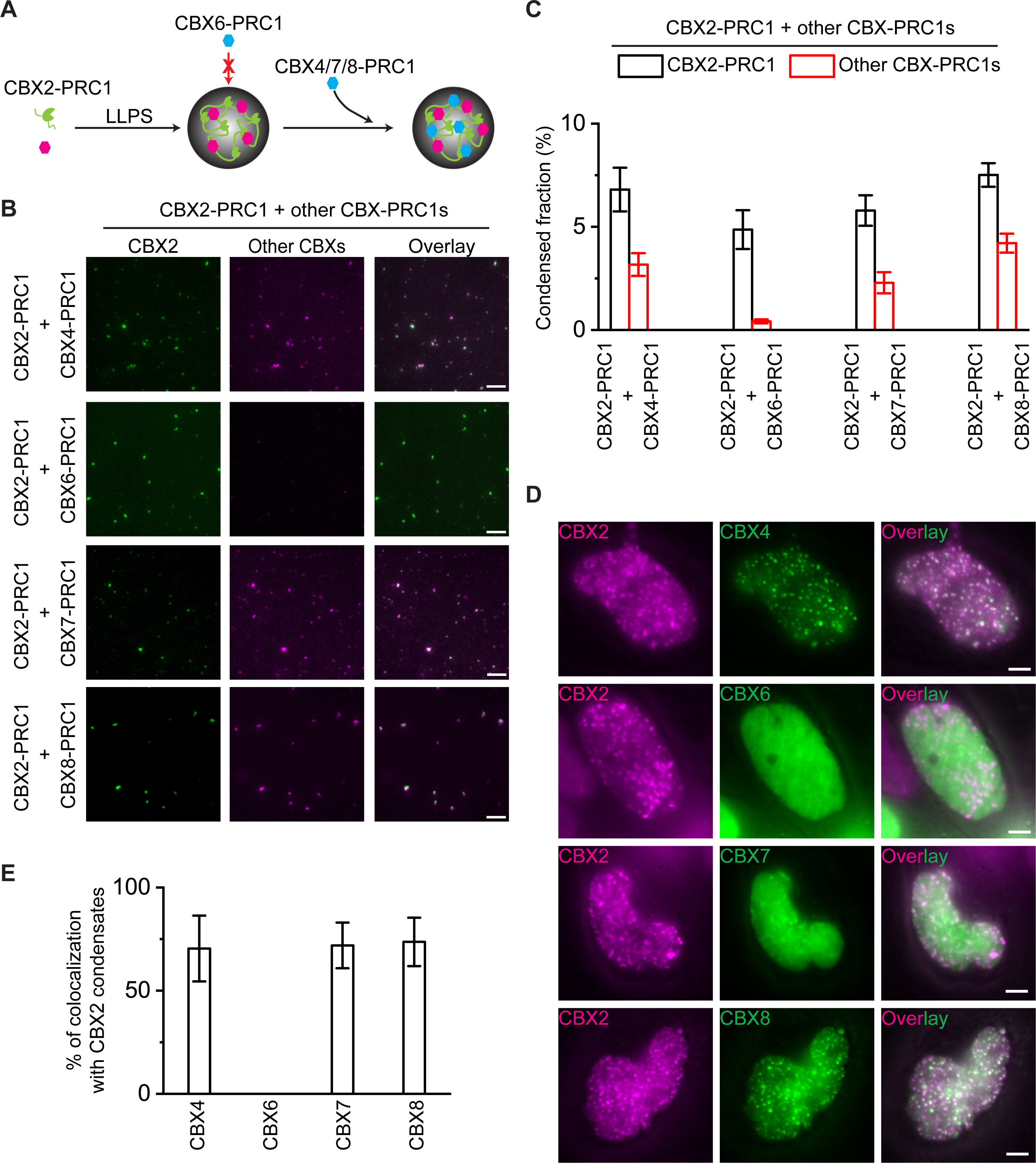
CBX2-PRC1 condensates recruit CBX4/7/8-PRC1 complexes but have a limited ability to condense CBX6-PRC1 in a compostion-dependent manner. (**A**) A hypothetical model describing how the CBX-PRC1 complexes are assembled into condensates. CBX2-PRC1 is assembled into condensates through a scaffold-client model. The CBX2-PRC1 condensates recruit CBX4/7/8-PRC1 but have a limited ability to enrich CBX6-PRC1. (**B**) Representative epi-fluorescence images of condensates of CBX2 and other CBX proteins (CBX4/6/7/8). CBX2-PRC1 (CBX2, RING1B, MEL18, and PHC1) was mixed with CBX4/6/7/8-PRC1 (CBX4/6/7/8, RING1B, MEL18, and PHC1), respectively. CBX2 was labelled with YFP while other CBXs were labelled with mCherry. Other CBX-PRC1 subunits (RING1B, MEL18, and PHC1) were not labelled. The concentration of individual components is 0.5 µM. Scale bar, 5.0 μm. (**C**) Condensed fraction of CBX2 and other CBX proteins quantified from Figure 5B. Data shown is from at least ten frames. Error bars are SD. (**D**) Live-cell epi-fluorescence images of HaloTag-CBX2 (HT-CBX2) and YFP-CBX4/6/7/8 in HeLa cells. HT-CBX2 fusion was stably integrated into the genome of HeLa cells. YFP-CBX4/6/7/8 were transiently expressed in HT-CBX2/HeLa cells, respectively. HT-CBX2 was labelled with HaloTag® TMR ligand. Scale bar, 5.0 µm. (**E**) Percentage of CBX4/6/7/8 condensates colocalization with CBX2 condensates quantified from Figure 5D. Data shown is from at least five cells. Error bars are SD.

To test whether these *in vitro* observations can be recapitulated in living cells, we transiently expressed YFP-CBX4/6/7/8 proteins in HeLa cells of which HT-CBX2 was stably integrated into the genome. After HT-CBX2 was labelled by HaloTag TMR ligand, live-cell epi-fluorescence imaging showed that the CBX2/4/7/8 proteins appear as condensates but as expected, CBX6 does not form condensates (**Figure 5D**). We then identified and segmented condensates of CBX2/4/7/8 and performed colocalization analysis of CBX2 condensates with the condensates of CBX4/7/8. We found that approximately 70% of condensates of CBX4/7/8 colocalize with condensates of CBX2 (**Figure 5E**). Thus, our cellular results support that CBX2-PRC1 condensates selectively recruit CBX4/7/8-PRC1s but cannot enrich CBX6-PRC1.

### Coarse-grained molecular dynamics simulations provide the molecular basis underpinning the scaffold-client model and its regulation

We have experimentally provided a concentration- and composition-dependent scaffold-client model that underpins the assembly of multi-component CBX-PRC1 condensates. In the following, we employed a multiscale molecular dynamics simulation approach to provide the molecular basis underpinning the scaffold-client model and regulation by the client RING1B. We first developed initial atomistic models of CBX2 and RING1B based on predictions from AlphaFold, to distinguish the protein regions that exhibit secondary structure from those that are expected to remain disordered (**Figure 6A**). We then developed coarse-grained representations using our one-bead per amino acid model (Joseph et al., 2021), which accounts for the charge, size, and relative hydrophobicity of each amino acid, and the secondary structure or disordered nature of the region they are in. With these coarse-grained residue-resolution models, we performed direct-coexistence simulations, where the dilute phase is simulated in coexistence with the condensate inside the same rectangular simulation box. From these simulations we estimated the stability of single-component condensates (of either CBX2 or RING1B) in the presence and absence of 10 mM Mg^2+^. By performing the direct-coexistence simulations for each single component system at various temperatures, we can estimate the coexistence densities of the dilute and dense phase (i.e., the condensate) as a function of temperature, and construct phase diagrams in the temperature-vs-density plane. From each phase diagram, we can obtain an estimate for the upper critical solution temperature, *T*_c_ (**Figure 6B**). Our residue-resolution phase diagrams revealed that while phase separation does not occur in the absence of Mg^2+^ ions and the absence of crowders (as evident from the low *T*_c_ of both systems), the ability of CBX2 and RING1B to undergo phase separation is significantly enhanced by the addition of 10 mM Mg^2+^ ions, and this increase in *T*_c_ occurs to a greater extent for CBX2.

**Figure 6.**
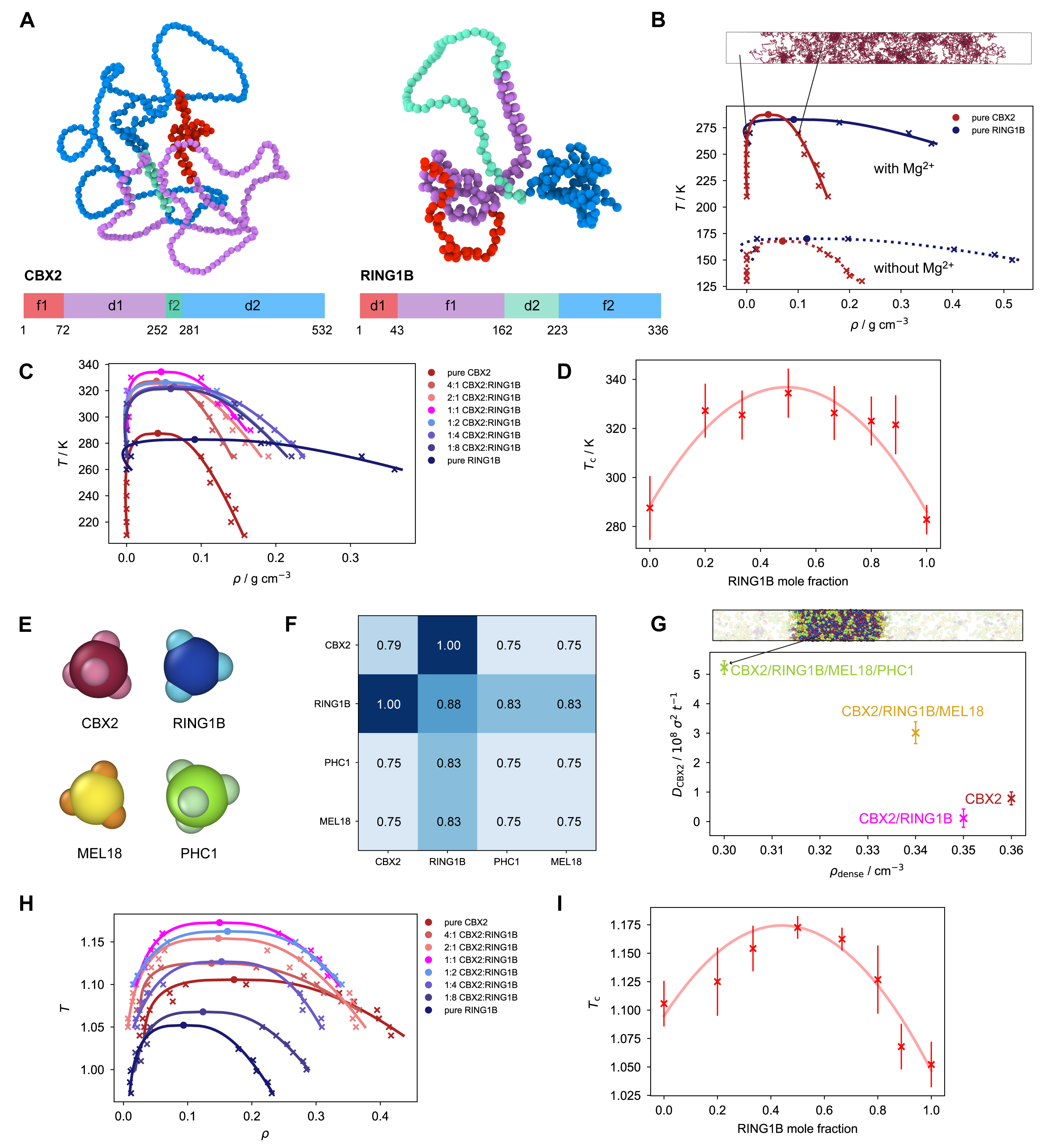
Coarse-grained molecular dynamics simulations provide the molecular basis underpinning the scaffold-client model and its regulation. (**A**) Residue-resolution coarse-grained model of CBX2 and RING1B. Residues in the folded regions are highlighted in red in the snapshots. In the schematic of the sequence, the different regions of the two proteins are referred to as ’f’ for folded and ’d’ for disordered and are colored in red and blue respectively. (**B**) Temperature(*T*)–density(*ρ*x) phase diagrams of pure CBX2 and RING1B systems in the presence of 10 mM Mg^2+^ (solid lines) and without (dotted lines). The addition of 10 mM Mg^2+^ increases the upper critical solution temperature, *T*_c_, of both systems but to greater extent for CBX2. Simulation snapshot depicts a pure CBX2 condensate in a direct-coexistence simulation at 270 K. (**C**) Temperature(*T*)–density(*ρ*) phase diagrams of pure CBX2 and RING1B systems, as well as their mixtures with varying RING1B mole fraction, in the presence of 10 mM Mg^2+^. (**D**) Variation of *T*_c_ of the systems plotted in **(C)**. We observe re-entrant behavior as a function of RING1B mole fraction, with the highest *T*_c_ (which corresponds roughly to the most stable condensate) observed in an approximately equimolar mixture of CBX2 and RING1B (i.e., RING1B mole fraction = 0.5). The red line is a second-order fit to the data points, and is intended as a visual guide only. (**E**) Minimal coarse-grained model of CBX2, RING1B, MEL18 and PHC1 that describes the proteins as patchy colloids. CBX2 and PHC1 are represented as 4-valency patchy particles, while RING1B and MEL18 are represented as 3-valency patchy particles. (**F**) Relative pairwise interaction strengths between the patches on the four different proteins in the minimal model at reduced temperature *T* = 1. The value of 1 represents the maximum strength of interaction considered in our simulations. These interactions were set according to the relative ability of the proteins to undergo phase separate, and the impact on the stability and material properties of multi-component condensates. (**G**) Diffusion coefficients of CBX2 measured in direct-coexistence simulations of the dense phase of mixtures with different compositions at *T* = 1. Each mixture contains an overall equimolar concentration of each protein. Simulation snapshot depicts a four-component mixture containing an equimolar amount of CBX2, RING1B, MEL18 and PHC1. The diffusion coefficient of CBX2 in the pure condensate and two-component mixture with RING1B is more similar in magnitude, while addition of MEL18 and PHC1 increases the diffusion coefficient of CBX2 in the three-and four-component mixture. (**H**) Temperature(*T*)–density(*ρ*) phase diagrams of pure CBX2 and RING1B systems, as well as their mixtures with varying RING1B mole fraction, using the minimal patchy-particle model. (**I**) Variation of *T*_c_ of the systems plotted in **(H)**. Again, we observe re-entrant behavior as a function of RING1B mole fraction, with the highest *T*_c_ observed in an equimolar mixture of CBX2 and RING1B. The red line is a second-order fit to the data points, and is intended as a visual guide only.

Next, we performed direct-coexistence simulations to investigate the stability of two-component condensates obtained from mixtures of CBX2 and RING1B mixed at different stoichiometric CBX2:RING1B ratios (4:1, 2:1, 1:1, 1:2, 1:4 and 1:8) (**Figure 6C**). In these two-component condensates, CBX2 behaves as the scaffold and RING1B as its client. The phase diagrams obtained from these simulations confirmed our experimental observations of the reentrant behaviour as a function of RING1B concentrations. Adding a small or moderate amount of RING1B (at a ratio of 2:1 or 1:1 CBX2:RING1B) modestly increases the stability of CBX2 condensates (**Figure 6D**). Adding larger amounts of RING1B (at a ratio of 1:4 or 1:8 CBX2:RING1B) notably decreases the stability of CBX2 condensates (**Figure 6D**). We confirmed this by repeating these simulations with another residue-resolution coarse-grained model, along with taking the effect of Mg^2+^ in a more approximate manner, reproduced similar trends in the relative stabilities of the condensates with different RING1B mole fraction (**Supplementary Figure 6A, B**).

To further probe the molecular driving forces for the reentrant phase behavior of the two-component CBX2-RING1B condensates as a function of RING1B concentration, as well as the effect of additional components (MEL18 and PHC1) on the liquid-like properties of Polycomb condensates, we performed simulations using a minimal coarse-grained model where proteins are represented as patchy colloids (Espinosa *et al*., 2020). That is, proteins are described by pseudo hard-sphere (PHS) particles decorated with sticky patches that allow the minimal proteins to establish multivalent transient interactions (**Figure 6E**). To recapitulate the relative ability of Polycomb proteins to phase separate that we observed experimentally, we represent CBX2 and PHC1 proteins, which can undergo phase separation on their own, with a 4-valency patchy particle with patches arranged in a tetrahedral arrangement on the hard-sphere core. For RING1B and MEL18, which behave as clients and do not phase separate on their own, we used 3-valency patchy particles. The strength of pairwise interactions between the different proteins were set according to their relative ability to phase separate, whether reentrant behaviour was observed experimentally as a function of the concentration of the protein, and the impact on the stability and material properties of multi-component Polycomb condensates. For example, since CBX2 has a greater propensity to undergo phase separation compared to PHC1, we set the per-patch CBX2–CBX2 interactions to be stronger than those of the PHC1–PHC1 interactions (**Figure 6F**).

Using this minimal model, we performed direct-coexistence simulations of the following systems: (1) pure CBX2 condensates, (2) CBX2/RING1B condensates, (3) CBX2/RING1B/MEL18 condensates, and (4) CBX2/RING1B/MEL18/PHC1 condensates. With the balance of interaction strengths shown in **Figure 6F** set in the minimal model, we were able to recapitulate the increasing liquid-like properties of the multi-component condensates with the addition of MEL18 and PHC1 into the Polycomb condensates, as seen from the increased diffusion coefficient of CBX2 when MEL18 and PHC1 were added (**Figure 6G**). Our simulations hence further support that the CBX2–MEL18 and CBX2–PHC1 interactions are weaker than the CBX2–CBX2 and CBX2–RING1B interactions present in the pure CBX2 and two-component CBX2/RING1B condensates, and how composition can tune the liquid-like behavior of CBX2 in Polycomb condensates.

For the two-component CBX2/RING1B condensates, we also performed simulations of mixtures with different CBX2:RING1B ratios (4:1, 2:1, 1:1, 1:2, 1:4 and 1:8) using our minimal patchy-particle model and from there, predicted the phase diagrams and upper critical solution temperature, *T*_c_, of these systems (**Figure 6H**). The simulations using the minimal model were similarly able to reproduce the reentrant behavior as a function of RING1B mole fraction observed in experiments and in the residue-resolution simulations, with the most stable condensate being formed at approximately equimolar concentration of CBX2 and RING1B (**Figure 6I**). We found that the reentrant behaviour originates from the cross CBX2–RING1B interactions being sufficiently stronger than the homotypic RING1B–RING1B and even the scaffold CBX2–CBX2 interactions. Thus, the equimolar mixture is expected to give the most stable condensate, since the enthalpic gain for condensate formation is maximized from maximizing the number of CBX2–RING1B interactions over the CBX2–CBX2 interactions. However, as the RING1B concentration continues to increase past the equimolar ratio, the stronger CBX2–RING1B and CBX2–CBX2 interactions start to get replaced for weaker RING1B–RING1B interactions. Altogether, our simulations from the minimal model highlight that, at the molecular level, the relative interactions strengths between the different Polycomb proteins are important in controlling both the stability and liquid-like properties of the Polycomb condensates.

## DISCUSSION

Studies have uncovered that CBX2 drives the formation of CBX-PRC1 condensates (Kent *et al*., 2020; Plys *et al*., 2019; Tatavosian *et al*., 2019); however, the molecular mechanisms underlying the assembly remain enigmatic, partly due to the complexity of CBX-PRC1 complexes. Here, we use biochemical reconstitution and cellular imaging to uncover the condensate assembly principles and utilize molecular dynamics simulations to supply a molecular basis for assembly. The main results and conclusions are summarized in **Figure 7**. CBX2 is the scaffold that drives the formation of CBX2-PRC1 condensates. The CBX2-PRC1 subunit clients regulate the reentrant phase transitions of CBX2 condensates, and the composition directs the partitioning and liquid-like behavior of CBX2-PRC1 condensates. CBX4/6/7/8 proteins show distinct abilities to form condensates. PRC1 subunits promote the condensation of CBX4 but dissolve CBX6/7/8 condensates. Finally, CBX2-PRC1 condensates selectively recruit CBX4/7/8-PRC1 but specifically exclude CBX6-PRC1. These results together suggest a context-dependent assembly of Polycomb condensates through a concentration- and composition-dependent scaffold-client model where CBX2 acts as a scaffold and the PRC1 subunits along with the other CBX proteins behave as clients.

**Figure 7.**
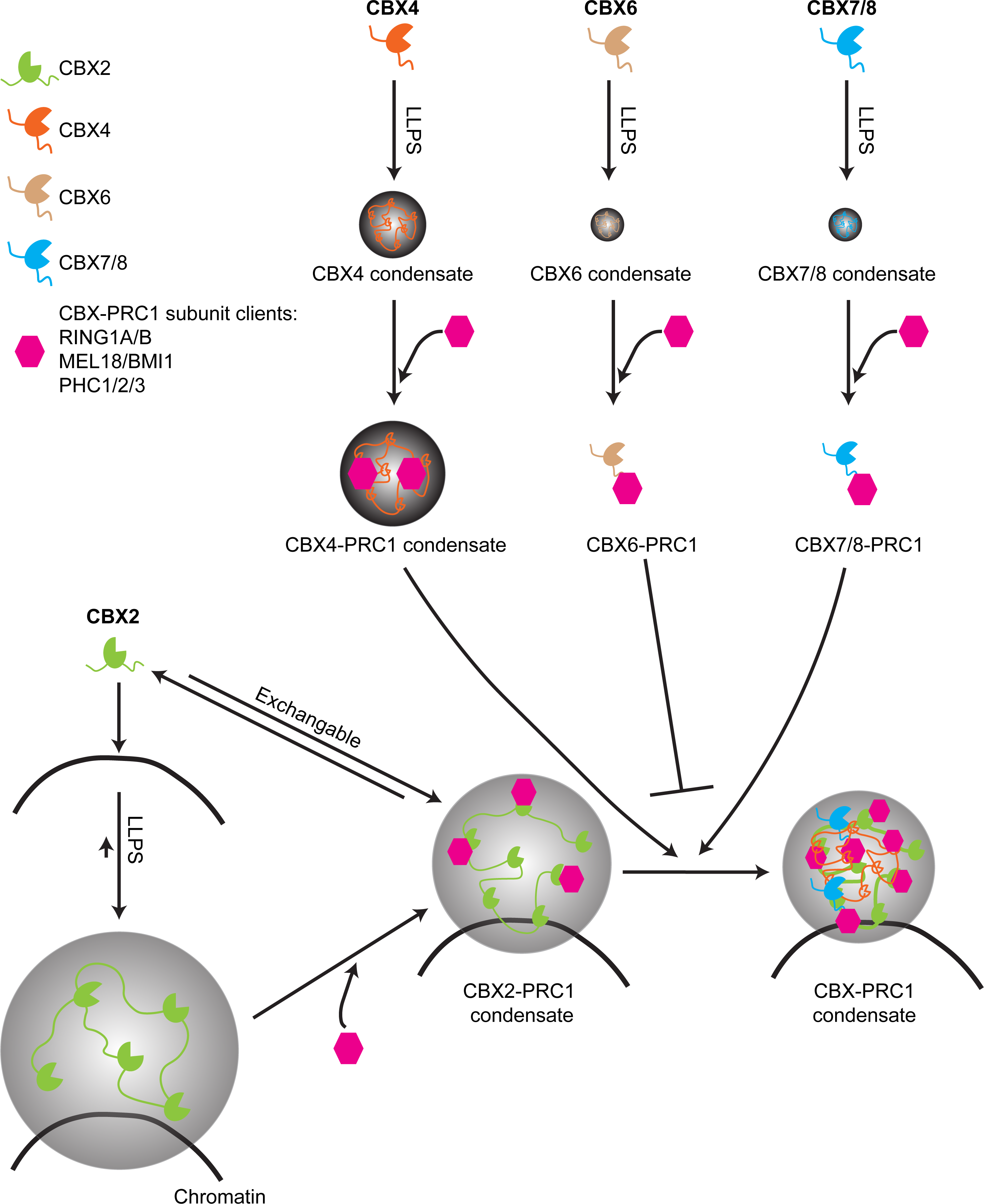
A model describing the assembly of CBX-PRC1 condensates. A proposed scaffold-client mechanism that underpins the biogenesis of CBX-PRC1 condensates through LLPS in a concentration- and composition-dependent manner. CBX2 seeds on chromatin and undergoes LLPS to form condensates that have a limited fluidity. The CBX2 condensates recruit the CBX-PRC1 subunit clients that regulate their partition in a concentration-dependent manner. The composition of CBX2-PRC1 condensates directs the fluidity of CBX2 and nucleosomes and regulates their partition. CBX4/6/7/8 form condensates with distinct abilities. CBX4 condensates recruit CBX-PRC1 subunit clients that in turn promote the assembly of CBX4-PRC1 condensates. The CBX-PRC1 subunit clients dissolve condensates of CBX6/7/8. CBX2-PRC1 condensates selectively recruit CBX4-PRC1 condensates and CBX7/8-PRC1 but have a limited ability to condense CBX6-PRC1. Given that CBX4-PRC1 can form condensates, it is possible that in the absence of CBX2-PRC1, CBX4-PRC1 can act as a scaffold. Our model implies a sequential event for the sake of simplicity and brevity; however, it should be noted that the order of events may be different from the proposed model.

### Scaffold-client assembly of CBX-PRC1 condensates

Our data show that the formation of CBX2-PRC1 multi-component condensates can be explained by the scaffold-client model of Banani et al.(Banani *et al*., 2016) The model postulates that within multi-component protein mixtures, just a few of the components, termed scaffolds, are essential to drive the system to phase separate. The enthalpic gain stemming from the formation of a liquid network of scaffold–scaffold interactions is sufficient to compensate for the entropic loss that occurs upon phase separation (i.e., condensates are associated with fewer microstates than the dissolved state). The other molecular components that are recruited to the condensates via their interactions with the scaffolds termed clients (Banani *et al*., 2016; Ditlev *et al*., 2018). While clients do not drive phase separation, they can sensitively modulate the properties of condensates (e.g., density and material properties). The scaffold-client model was initially developed based on simple reconstituted systems, where the assembly of condensates is regulated by the valence and concentration of scaffold molecules (Banani *et al*., 2016). More recently, the scaffold-client model has been exemplified by the assembly of stress granules where G3BP1 drives condensate formation (Guillen-Boixet et al., 2020; Sanders et al., 2020; Yang et al., 2020) and is further supported by simulations (Espinosa *et al*., 2020).

A few lines of evidence support that CBX2 is the scaffold and drives the formation of the multi-component CBX2-PRC1 condensates under the conditions tested: (1) Among the components of CBX2-PRC1, CBX2 demixes at concentrations as low as 10 nM, in the presence of low concentrations of Mg2+ ions and absence of crowders. Whereas other components do not form condensates or form condensates at much higher concentrations than CBX2. (2) CBX2 condensates enrich other CBX2-PRC1 components. (3) In living cells, increasing the CBX2 concentration increases its condensed fraction; however, increasing other CBX2-PRC1 subunits decreases their condensed fraction. (4) Among the CBX proteins, CBX2 has the best ability to condense. (5) Increasing the concentration of other CBX proteins in living cells does not increase their condensed fraction. (6) CBX2-PRC1 condensates enrich CBX4/7/8-PRC1s but exclude CBX6-PRC1. (7) Coarse-grained molecular dynamics simulations show that CBX2 condensates are more stable than RING1B. Overall, our results supply compelling evidence by which CBX2 is the scaffold of CBX-PRC1 condensates. Unlike CBX6-, CBX7-, and CBX8-PRC1, CBX4-PRC1 can form condensates independently of CBX2-PRC1. Since the expression of CBX proteins is developmentally regulated (Kim and Kingston, 2022; Piunti and Shilatifard, 2021), it is possible that in the absence of CBX2, CBX4 may organize CBX-PRC1 condensates. However, in mES cells, CBX2 should be the scaffold since CBX4 is not expressed (Camahort and Cowan, 2012). Further studies are needed to address how the assembly of CBX-PRC1 condensates is regulated during development and differentiation.

Our results show that the PHC proteins can undergo LLPS and their LLPS ability is not due to polymerization, which is consistent with earlier reports (Seif *et al*., 2020). However, the condensation ability of PHC proteins is much less efficient than CBX2 and CBX4. Both CBX2 and CBX4 can enrich PHC1 within their condensates. It is interesting to note that the inclusion of PHC1 in the multi-component CBX2-PRC1 complex inhibits CBX2 condensate formation, while adding PHC1 in the multi-component CBX4-PRC1 promotes CBX4 condensate formation. In some cellular contexts, overexpression of the PHC proteins can promote the Polycomb condensate assembly (Isono *et al*., 2013). Furthermore, the inclusion of PHC1 dissolves condensates of CBX6/7/8. As such, these results suggest a CBX-specific role of the PHC proteins in the assembly of CBX-PRC1 condensates.

### Stoichiometry- and composition-dependent control of CBX-PRC1 condensate assembly

The formation and dissolution of phase-separated condensates is essential for regulating condensate-directed functions. Here we show that the relative concentration of the scaffold CBX2 to the client RING1B controls the assembly and disassembly of heterotypic CBX2-RING1B condensates: a relatively high ratio of CBX2 to RING1B promotes condensate formation while a relatively low ratio of CBX2 to RING1B dissolves condensates. This phase behavior is termed as a reentrant phase transition (Milin and Deniz, 2018). Our simulations show that this reentrant phase transition of CBX2 occur because CBX2 and RING1B bind to one another much more strongly than the CBX2–CBX2 self-interactions. The reentrant phase behavior seen for CBX2 is much like the RNA-driven reentrant phase behaviour seen for FUS, where FUS phase separation is stabilized by addition of small concentrations of RNA but abrogated at high RNA concentrations (Banerjee *et al*., 2017). The reentrant phase transitions supply a potential mechanism for controlling transcriptional condensate assembly through RNA (Henninger *et al*., 2021). The stoichiometry-dependent assembly/disassembly orchestrates actomyosin cortex activation (Yan et al., 2022). The protein level of RING1B in mES cells is much higher than CBX2, suggesting that RING1B can dissolve CBX2 condensates. Consistent with this, CBX2 shows an increased condensed fraction and a larger condensate size in *Ring1a^-/-^/1b^-/-^* mES cells when compared to wild type mES cells. As such, these results show a concentration-dependent assembly of CBX2-PRC1 condensates. The expression level of Polycomb components changes during development and tumorigenesis (Kim and Kingston, 2022). In the future, it is important to dissect how variations in expression are linked with the assembly of condensates that contribute to physiology and pathology.

Our results show that the composition of heterotypic multi-component CBX2-PRC1 system directs the assembly and biophysical properties of condensates. The inclusion of more components in the system dissolves condensates and excludes nucleosomes but increases the liquid-like behavior of CBX2 and nucleosomes. These results suggest that the assembly and biophysical properties of CBX-PRC1 condensates are regulated by their composition without a fixed saturation concentration. Consistent with this, recent studies demonstrate using model systems of biological condensates that composition-dependent heterotypic LLPS drives condensate assembly, subunit exclusion (Riback *et al*., 2020), and the dynamics of components (Ferrolino *et al*., 2018).

The composition-dependent control of partitioning and dynamics of CBX2 and nucleosomes could be important for controlling chromatin states during cell cycle or cell differentiation. Our earlier studies have shown that CBX2 is fluid in the interphase but is gel/solid during the metaphase (Zhen et al., 2014). During mES cell differentiation, the dynamics of CBX2 progressively becomes slow (Ren *et al*., 2008). On the other hand, chromatin is highly dynamic in mES cells and becomes less dynamic during differentiation (Meshorer et al., 2006). It is possible that the composition-dependent control of assembly and properties of CBX-PRC1 condensates couples with the dynamics of CBX2 and nucleosomes during cell-fate transitions and cell cycle progression, linking with epigenetic inheritance and establishment/maintenance.

### Implications of the scaffold-client assembly for genome organization

Both PRC2 and PRC1 are involved in the long-range clustering of Polycomb target genes (Schuettengruber *et al*., 2017). Here, we intend to connect earlier observations by using our scaffold-client assembly of CBX-PRC1 condensates and build a testable model for future investigations. We utilize mES cells to illustrate our model for the simplicity of CBX-PRC1 complexes in mES cells since CBX4 and CBX8 do not express in mES cells and CBX6 does not form condensates. As such, we only consider CBX2-PRC1 and CBX7-PRC1 in mES cells. We propose the following sequential events to unify earlier observations. CBX2 seeds onto chromatin through direct interactions with underlying DNA sequences to assemble CBX2-PRC1 condensates. It has shown that CBX2 interacts with AT-rich DNA, which is essential for the assembly of CBX2-PRC1 condensates (Kent *et al*., 2020). CBX2-PRC1 condensates compact chromatin locally, consistent with the observations that CBX2-PRC1 condensates enrich nucleosomes. CBX7-PRC1 acts as a client which brings distal H3K27me3-marked chromatin into CBX2-PRC1 condensates. The polymerization of PHC family proteins may help recruiting CBX7-PRC1 bound H3K27me3-chromatin into CBX2 condensates. The removal of H3K27me3 and the subsequent disruption of polymerization can impair the recruitment of H3K27me3-chromatin into CBX2-condensates for compaction. The disruption of RING1B will impair the CBX7-PRC1 complex as well as the assembly of CBX2-PRC1 condensates. As such, our current results pave the way to dissect the molecular mechanisms underlying the Polycomb-mediated genome organization through LLPS.

Overall, our studies uncover the molecular principles underlying the assembly of CBX-PRC1 condensates through phase separation. We propose that the composition- and concentration-dependent scaffold-client assembly of multi-component CBX-PRC1 condensates is coupled with genome organization and links with epigenetic inheritance. As such, our studies supply the foundations for further exploring the functional roles of Polycomb condensates through LLPS.

## METHODS

### Cell lines

pGK12.1 mES cells (Penny et al., 1996) were provided by Dr. Neil Brockdoff (University of Oxford, UK). *HT-CBX2/Ring1a^−/−^/Ring1b^fl/fl^; Rosa26::CreERT2* mES cells (referred to as *HT-CBX2/Ring1a^-/-^/Ring1b^fl/fl^* thereafter) and *HT-CBX2/Mel18^−/−^/Bmi1^−/−^* mES cells were reported previously (Kent *et al*., 2020). pGK12.1 cells expressing *HT-CBX2* or *YFP-CBX2* were reported previously (Kent *et al*., 2020; Ren *et al*., 2008). *HT-PRC1* subunits were randomly integrated into the genome of HeLa cells as noted in the text. pGK12.1 cells carrying HaloTag (HT) knockin at the C-terminus of two *CBX2* alleles (homozygous, *Cbx2^HT/HT^*) and one *CBX2* allele (heterozygous, *CBX2^WT/^*^HT^) were generated as described in the text.

### Cell culture

pGK12.1 mES cells were grown in Dulbecco’s Modified Eagle Medium (Sigma-Aldrich; D5796,) supplemented with 15% Fetal Bovine Serum (VWR; 97068-085), 0.1 mM β-mercaptoethanol (Gibco; 31350-010), 2.0 mM L-Glutamine (Gibco; 25030149), 100 Units/mL Penicillin-Streptomycin (Gibco; 15140-122), 1 × MEM Non-Essential Amino Acids (Gibco; 11140-050), 10 µg/mL Ciprofloxacin (Sigma; 17850), and 10^3^ units/mL leukemia inhibitor factor (LIF) at 37 °C in 5% CO_2_. HEK293T cells were grown in DMEM supplemented with 10% FBS, 0.1 mM β-mercaptoethanol, 2.0 mM L-Glutamine, 100 Units/mL Penicillin-Streptomycin, and 10 µg/mL Ciprofloxacin (Sigma; 17850) at 37 °C in 5% CO_2_.

To deplete *Ring1b*, *Ring1a^−/−^*/*Ring1b^fl/fl^*; *Rosa26*::CreERT2 mES cells were treated with 1.0 µM 4-Hydroxytamoxifen (H7904, Sigma) for 2 days as described previously (Zhen *et al*., 2014).

### Plasmids

#### Plasmids for recombinant protein production

The pGEX-6P-1-GST plasmid has been described previously (Tatavosian *et al*., 2019) and contains an ampicillin resistance gene. To clone pGEX vectors with a fluorescence protein, we used polymerase chain reaction (PCR) to generate Cerulean-FLAG, YFP-FLAG, and mCherry-FLAG. This was ligated downstream of the sequence encoding glutathione S-transferase (GST) within pGEX-6P-1-GST. Once pGEX-GST-YFP-FLAG had been generated, PCR was used to amplify *CBX2,* and it was ligated between the sequences encoding the GST and YFP motifs. To generate pGEX-GST-CBX2-FLAG, an oligonucleotide encoding for FLAG was used to replace YFP-FLAG in pGEX-GST-CBX2-YFP-FLAG. Once pGEX-GST-mCherry-FLAG had been generated, PCR was used to amplify CBX4, CBX6, CBX7, CBX8, and MEL18 and one was added between the sequences encoding GST and mCherry to generate the pGEX-GST-CBX4/6/7/8-mCherry-FLAG and pGEX-GST-MEL18-mCherry-FLAG plasmids. To generate pGEX-GST-MEL18-YFP-FLAG, *MEL18* was cut from pGEX-GST-MEL18-mCherry-FLAG and ligated into pGEX-GST-YFP-FLAG between GST and YFP. Once pGEX-GST-Cerulean-FLAG had been generated, PCR was used to amplify PHC1 and RING1B. This was ligated between the sequences GST and Cerulean. Cerulean-FLAG in pGEX-GST-RING1B-Cerulean-FLAG was replaced with FLAG oligonucleotide to generate pGEX-GST-RING1B-FLAG. *CBX2* in pGEX-GST-CBX2-YFP-FLAG was replaced with *RING1B* to generate pGEX-GST-RING1B-YFP-FLAG. To generate pGEX-GST-MEL18-FLAG, RING1B was replaced with MEL18 in pGEX-GST-RING1B-FLAG. Oligonucleotides were created to change leucine 307 to arginine (CTC to CGC) within PHC1 to generate PHC1(L307R). The relevant section was removed from pGEX-GST-PHC1-Cerulean-FLAG and was then replaced with the PHC1(L307R) oligonucleotide to generate pGEX-GST-PHC1(L307R)-Cerulean-FLAG. A PreScission Protease recognition site was added between PHC1 and Cerulean-FLAG by oligonucleotide ligation to generate pGEX-GST-PHC1-Protease-Cerulean-FLAG. An oligonucleotide encoding the sequence of FLAG was ligated into the PHC1(L307R) plasmid to generate pGEX-GST-PHC1(L307R)-FLAG. To generate pGEX-GST-RING1A-Cerulean-FLAG, the sequence encoding RING1A was amplified by PCR. RING1B was removed from pGEX-GST-RING1B-Cerulean-FLAG and replaced with RING1A. To generate pGEX-6P-1-GST-BMI1-mCherry-FLAG, BMI1 was amplified using PCR. Then, MEL18 was removed from pGEX-GST-MEL18-mCherry FLAG and replaced with BMI1. To generate pGEX-6P-1-GST-PHC2-Cerulean-FLAG, PHC2 was amplified from pEZ-M39-PHC2-FLAG (EX-L3818-M39, Genecopoeia) and was ligated into pGEX-GST-Cerulean-FLAG. pGEX-GST-6P-1-GST-PHC3 was generated similarly with PCR amplification of PHC3 using pEZ-M39-PHC3-FLAG (EX-L3818-M39, Genecopoeia) and ligation into pGEX-GST-PHC2-Cerulean FLAG after digestion to remove PHC2. The sequences of plasmids were verified by Sanger sequencing.

#### Plasmids for expression in mammalian cells

Plasmids pTripZ-M1-FLAG-HT-CBX2/4/6/7/8 were reported previously (Zhen et al., 2016). pTripZ-M1-FLAG-HT-MEL18, pTripZ-M1-FLAG-HT-PHC1, and pTripZ-M1-FLAG-HT-RING1B have also been previously reported (Kent *et al*., 2020). These pTripZ plasmids have the ampicillin and zeocin resistance genes as well as the puromycin resistance gene. pTripZ-G2-YFP-CBX2(ΔTRE) was reported previously (Tatavosian *et al*., 2019) and contains ampicillin, zeocin, and geneticin resistances. To generate pTripZ-M1-FLAG-HT-BMI1 and pTripZ-M1-FLAG-HT-RING1A, BMI1 and RING1A were amplified by PCR and replaced CBX6 in pTripZ-M1-FLAG-HT-CBX6. pTripZ-M1-FLAG-HT-PHC2 was generated through PCR amplification of PHC2 from pEZ-M39-PHC2-FLAG and replacement CBX6 in pTripZ-M1-FLAG-HT-CBX6. pTripZ-M1-FLAG-HT-PHC3 was generated similarly using pEZ-M39-PHC3-FLAG as a template for PCR. These plasmids were used to randomly integrate HT-PRC1 subunit fusion proteins to the genome of HeLa cells.

### CRISPR/Cas9-mediated HDR

To endogenously tag the C-terminus of *CBX2* with HaloTag (HT), a sgRNA was designed using the IDT’s ‘design custom gRNA’ tool, and sgRNA oligos synthesized from IDT were annealed and cloned into pSpCas9(BB)-2A-Puro (PX459)-V2.0 (Addgene; 62988). Homology arms for the donor vector were amplified via PCR from digested DNA fragments of genomic DNA purified from mES cells and cloned into a premade pUC19-HT vector. Approximately 2 million mES cells were mixed with 25 µg donor vector and 15 µg pX459-sgRNA and incubated on ice for 10 minutes. Electroporation was performed using a Gene Pulser XCell in 600 µL Ingenio electroporation buffer (Mirus Bio; MIR 50114) with the settings of 260V & 500 μF using an exponential decay in a 0.4 cm cuvette (Bio-Rad; 165-2088). After electroporation, cells were incubated on ice for an additional 10 minutes before added to warm medium and plated on a gelatinized plate. The morning after electroporation, 1.0 µg/ml puromycin was added to the cell medium for 48 hours to select for cells that received the sgRNA vector. After approximately 1-week, colonies were picked and then expanded. A portion of cells were labeled with 100 nM HaloTag TMR ligand (Promega; G8252) and fluorescent colonies were identified using a TIRF microscope. Genomic DNA from fluorescent colonies was purified, and proper editing of the genome was verified via PCR. Colonies that had either a homozygous or heterozygous insertion were further screened with a western blot using Anti-HaloTag (Promega; G921A) and compared with the parental cell line as a negative control for final validation.

#### CBX2-C-Tag sgRNA Oligos

sgRNA-CBX2-C-TOP: CACCGCAAGTTGAAGAAGCCCACGC

sgRNA-CBX2-C-BOT: AAACGCGTGGGCTTCTTCAACTTGC

#### CBX2 Homology Arm PCR Amplification Primers

CBX2-C-LArm-Fwd: ATGATTACGCCAAGCTTGCATGCCTGCAGGTCGACTCTAGACACGTGAAAGACCACCAGGAAGGATCTGGGGA

CBX2-C-LArm-Rev: TTCACGCGTGCTATAATGCCTCAAGTTGAAGAAGCCCACGCTAGTGGGCGACTCCTT

CBX2-C-RArm-Fwd: AGCGGCCGCCAGACAATTGCCCGAGCCAGACCTGCCTT CBX2-C-RArm-Rev:

ACGACGGCCAGTGAATTCGAGCTCGGTACCCGGGGATCCTACGTATCACCAGGCGGAGCTCTGCACCA

#### CBX2-C-Tag PCR Verification Primers

CRISPRverify-CBX2-C-Fwd: AAACGGGGACGCAAGCCTCTAC

CRISPRverify-CBX2-C-Rev: GCTCCTAAACGACCCCACCACAC

### Establishing stable cell lines

To generate stable HeLa cell lines, pTripZ-M1 plasmids containing PRC1 subunit fusion gene and the puromycin selective marker as well as the pTripZ-G2-YFP-CBX2(ΔTRE) plasmid harboring a geneticin selective marker were purified according to the protocol listed in EndoFree Max Plasmid Kit (QIAGEN; 12362). 24-hours before transfection, human embryonic kidney cells with the T7 antigen (HEK293T) were seeded in a 100-mm dish to reach 85-95% confluency at the time of transfection. Cells were co-transfected with 21 µg pTripZ, 21 µg psPAX2, and 10.5 µg pMD2.G by using calcium phosphate precipitation as described previously (Tatavosian et al., 2015; Zhen *et al*., 2014). 11 hours after transfection, the medium was changed with 10 mL fresh culture medium and cells were incubated for 48 hours at 37°C, 5% CO_2_. Viral titer in medium was collected and cell debris was pelleted by centrifugation at 1,000 × g for 5 minutes at 4°C. A single-cell suspension of HeLa cells was obtained from an 80% confluent 100-mm dish and 30% of this was transferred to a 15 mL new tube and collected via centrifugation at 700 × g for 5 minutes at 4°C. HeLa cells were added to virus-containing medium supplemented with 1.0 mL of 5 × culture medium and 8 µg/mL polybrene. For dual-infection cell lines, 2.0 mL of 5 × culture medium was added to the combined viral titer from two relevant transfections. The solution was plated in a 100-mm dish prepared with 0.2% gelatin and incubated at 37°C, 5% CO_2_ for 12 hours. Cells were washed once with PBS solution (Sigma, D8537) and 12 mL fresh culture medium was added. Cells were grown for 72 hours, or until plate reached 50% confluency. Puromycin was added at 2 µg/mL to select stably integrated cells for at least one week, and culture was continued with 1 µg/mL puromycin afterwards. All dual infections contained one plasmid that conferred puromycin resistance and another that conferred geneticin resistance. Thus, puromycin was added to 2 µg/mL and geneticin was added to 600 ng/mL for at least one week to perform selection in these cell lines.

### Generating recombinant protein

The expression vector pGEX-6P-1-GST-FLAG containing the relevant fusion protein and fluorescence protein was transformed into Rosetta™ 2 (pLysS) host strains (Novagen; 71403). A single colony was inoculated in 5.5 mL LB broth (Fisher Scientific; BP1426) with 100 µg/mL ampicillin at 37°C for 16-18 hours while shaking at 250 rpm. The following morning, 4.0 mL of overnight culture was transferred to 1.0 L of fresh LB and shook at 250 rpm, 37°C for 7-8 hours to reach OD_600_ ≥1.5. To induce protein expression, 1.0 mM IPTG (IBI Scientific; IB02105) was added to the culture and the culture was shaken at 18°C for 16-18 hours. *E. Coli* was harvested via centrifugation at 5,000 × g for 15 minutes at 4°C and resuspended in 25 mL lysis buffer (50 mM HEPES pH 7.5, 0.5 mM EDTA, 1.6 M KCl, 0.5 mM MgCl_2_, 1 mg/mL lysozyme, 1 mM DTT, 0.2 mM PMSF, 20 µg/mL RNase A, and 1 × protease inhibitor cocktail (Sigma; S8830)). Three freeze-thaws cycling between -80°C and an ice-water bath were used to partially lyse bacteria. Cells were completely lysed via sonication (Vibra-Cell™; VCX130) for 7.0 minutes using 65% amplitude and cycles at 15 seconds on-time, 45 seconds off-time (28 cycles total). IGEPAL^®^ CA-630 was added to 0.1% and the solution was rocked at 4°C for 30 minutes before centrifugation at 15,000 × g for 15 minutes to pellet cell debris. To precipitate DNA and RNA, PEI (Poly(ethyleneimine), Sigma-Aldrich; P3143) was added dropwise to 0.30% while solution was vortexed and then the solution was rocked for 10 minutes at 4°C before DNA/RNA was collected by centrifugation at 20,000 × g for 10 minutes at 4°C. To ensure complete DNA/RNA precipitation, PEI was added dropwise to 0.30% and solution was rocked for 20 minutes at 4°C and DNA/RNA was collected by centrifugation at 20,000 × g for 10 minutes. Supernatant was added to 750 µL Pierce™ Glutathione Agarose (Thermo Scientific; 16100) pre-washed with cold 1× PBS (Sigma-Aldrich; P4417) and rocked for 1 hour at 4°C while protected from light. Beads were collected via centrifugation at 700 × g for 2 minutes at 4°C and supernatant was carefully removed. Beads were washed with 8-10 mL chilled Buffer A (20 mM HEPES pH 7.5, 0.2 mM EDTA, 1.0 M KCl, 0.2 mM PMSF, and 1 mM DTT) and collected by centrifugation at 700 × g at 4°C for 2 minutes. Supernatant was removed and beads were transferred to a Poly-Prep Chromatography Column (Bio-Rad; 731-1550) where they were washed twice with 10 mL chilled Buffer A and allowed to drain by gravity. To elute recombinant protein from Pierce agarose beads, 500 µL of GSH Elution Buffer pH 8.0 (Buffer A + 80 mM reduced L-Glutathione (Sigma-Aldrich; G4251) + Tris-Base) was incubated for 10 minutes and then collected. Elution was repeated with 3 × 500 µL additions to reach 1.5 mL total volume. Collected protein was incubated 1 hour with 200 µL ANTI-FLAG-M2 affinity resin (Sigma-Aldrich; A2220) pre-washed with 3 times with1.0 mL Buffer B (20 mM HEPES pH 7.5, 0.2 mM EDTA, 1.0 M KCl, 0.2 mM PMSF, and 1 mM DTT). Affinity resin was collected via centrifugation at 500 × g at 4°C for 2 minutes. Resin was washed a total of three times with 1.0 mL Buffer B and collected by centrifugation to remove supernatant. To elute protein, ANTI-FLAG resin was rocked with 300 µL Buffer BC (20 mM HEPES pH 7.5, 0.2 mM EDTA, 0.05% IGEPAL^®^ CA-630, 20% glycerol, 0.5 mM DTT, 0.1 mM PMSF, 1 × protease inhibitor cocktail (Sigma-Aldrich; P1860), 300 mM KCl, and 0.8 mg/mL FLAG peptide (Sigma-Aldrich; F3290) at 4°C for 1 hour. Resin was collected via centrifugation at 2,000 × g for 2 minutes at 4°C. Supernatant was collected and transferred to a new tube before being spun at 20,000 × g for 10 minutes. This supernatant was collected and transferred to a 0.5 mL Pierce™ concentrator, PES (Thermo Scientific; 88513) where it was concentrated down to 20-30 µL by centrifugation at 15,000 × g for 45-50 minutes at 4°C in a fixed-angle rotor. Recombinant protein was resolved by NuPAGE™ 4-12% Bis-Tris Gel (Invitrogen; NP0322BOX) to determine identity and purity. Protein concentration was quantified by measuring absorption at 595 nm using the Pierce™ Detergent Compatible Bradford Assay Reagent (Thermo Scientific;1863028).

### Nucleosome reconstitution

To generate 601 (Widom) DNAs used for Cy3-labeled nucleosome reconstitution, pGEM-3z 601 (Addgene; 26656) was amplified by PCR with a Cy3-labeled forward primer. PCR amplicon was purified twice using PCR purification kit (QIAGEN; 28104). Fragment size was confirmed using agarose gel electrophoresis and the DNA concentration was quantified using absorbance at 547 nm. Histone octamer, recombinant human with H3 K_c_27me3, was purchased from the Histone Source, Colorado State University (HOCT-H3K27me3-0.5mg). The reconstitution was performed as described in (Luger et al., 1999). 1.0 µM 601 DNA and 1.0 µM histone octamer were mixed in 10 µL of 10 × Nucleosome Reconstitution Buffer (NRB; 100 mM Tris-HCl, pH 7.4, 10 mM DTT, 10 mM EDTA, and 1.0 mg/mL BSA) containing 2.0 M NaCl. Aliquots of 1 × NRB were added every 12-15 minutes to achieve a final nucleosome concentration at 10 nM. The reconstituted nucleosomes are stored in -80°C until use.

### Optical setup for epi-fluorescent microscopy

Epi-fluorescent images were captured using a Zeiss Axio Observer D1 Microscopy (Zeiss, Germany) equipped with an Evolve 512 × 512 EMCCD camera with pixel size 16 µm (Photometrics; Tucson, AZ). For *in vitro* imaging of condensates, an Alpha Plan-Apochromatic 100×/1.40 NA Oil-immersion Objective (Zeiss, Germany) was used. For the excitation and emission of YFP, a Brightline^®^ single-band laser filter set (Semrock; excitation filter, FF02-482/18-25; emission filter, FF01-525/25-25; dichroic mirror, Di02-R488-25) was used. For Cerulean fluorescence, a Brightline^®^ single-band laser filter set (Semrock; excitation, FF02-438/24-25; emission filter, FF01-483/32-25) was used. For mCherry fluorescence, 560/10-nm excitation and 610/35-nm emission filters were used. For imaging of live-cell condensates, an Alpha Plan-Apochromatic 100×/1.46 NA Oil-immersion Objective (Zeiss, Germany) was used. For the excitation and emission of TMR, a Brightline® single-band laser filter set (Semrock; excitation filter, FF01-561/14; emission filter, FF01-609/54; dichroic mirror, Di02-R561-25) was used. For imaging of immunostained cells, an Alpha Plan-Apochromatic 100×/1.46 NA Oil-immersion Objective (Zeiss, Germany) was used. For the excitation and emission of Alexa 488 nm, a Brightline^®^ single-band laser filter set (Semrock; excitation filter, FF02-482/18-25; emission filter, FF01-525/25-25; dichroic mirror, Di02-R488-25) was used. For Alexa 568 nm fluorescence, 560/10-nm excitation and 610/35-nm emission filters were used. The microscope and the EMCCD camera were controlled by Slidebook 6.0 software (Intelligent Imaging Innovations, Colorado).

### *In vitro* condensate formation

For investigating the condensation ability of individual subunits of CBX-PRC1, purified proteins were serially diluted in the condensation formation buffer (20 mM HEPES pH 7.5, 1.0 mM MgCl_2_, and 100 mM KCl). For interrogating the recruiting of PRC1 subunits into CBX2 condensates, CBX2-YFP was fixed at 0.5 µM in the condensation formation buffer while varying concentrations of RING1B-Cerulean, MEL18-mCherry, and PHC1-Cerulean, respectively. For studying the recruiting of CBX proteins to CBX2 condensates, 0.5 µM of CBX2-YFP was mixed with 0.5 µM of CBX4-mCherry, CBX6-mCherry, CBX7-mCherry, and CBX8-mCherry, respectively, in the condensation formation buffer. For studying the recruiting of CBX proteins into CBX2-PRC1 condensates, CBX4/6/7/8-mCherry proteins were, respectively, added into the mixture of 0.5 µM of each CBX2-YFP, RING1B, PHC1, and MEL18 in the condensate formation buffer. To test the effect of the complex composition on the condensate formation for each CBX2-PRC1 subunit, different preparations of 3 fluorophores were prepared: CBX2-YFP, MEL18-mCherry, RING1B, and PHC1-Cerulean; or CBX2, MEL18-mCherry, RING1B-YFP, and PHC1-Cerulean, with 0.5 µM of each component. To test the effect of the complex composition on the condensate formation of each CBX4-PRC1 subunits, different preparations of 3 fluorophores were prepared: CBX4-mCherry, MEL18, RING1B-YFP, and PHC1-Cerulean; or CBX4-mCherry, MEL18-YFP, RING1B, and PHC1-Cerulean, with 0.5 µM of each component. These same experiments were repeated for CBX6-mCherry, CBX7-mCherry, and CBX8-mCherry.

Unless otherwise indicated, the solution was incubated for 20 minutes at room temperature after being mixed. The 10 µL sample was placed in an 8-9 mm × 1.7 mm CoverWell™ Perfusion Chamber (Grace Bio-Labs; 622205) adhered to a clean 25 × 25 mm micro cover glass (VWR; 48366249) and the holes were covered with tape to prevent sample evaporation. Images were taken of condensates that settled to the surface of the cover glass using the imaging configuration listed in “Optical setup for epi-fluorescent microscopy” section. Images are presented by Photoshop and ImageJ. Images were analyzed by using CellProfiler as described in “Quantifying condensed fraction and condensate size, and colocalization by CellProfiler” section.

### Live-cell imaging of condensates

Cells containing HT-PRC1 fusion gene were incubated without or with 0.4 µg/mL doxycycline hyclate (Sigma-Aldrich; D9891) for 72 hours prior to imaging. A 35-mm cover-glass-bottom dish was coated with gelatin the day before splitting cells. The next day, HeLa cells containing PRC1 fusion gene were plated evenly on coated 35-mm dish and incubated overnight at 37°C, 5% CO_2_. To visualize condensate formation, 50 nM HaloTag TMR ligand was added to culture medium of plates and incubated for 15 minutes at 37°C, 5% CO_2_. Medium was removed and plate was washed once with fresh culture medium. A new medium was added, and the plate was incubated for 30 minutes at 37°C, 5% CO_2_. The recovery process was repeated for a total of one hour of recovery after dye incubation. The plate was washed once with PBS and then loaded with imaging medium (Life Technologies; A1896701) supplemented with 10% FBS and 10^3^ units/mL LIF. Cells were imaged for less than 90 minutes while temperature was maintained at 37 °C using a heater controller (Warner Instrument; TC-324). Images were acquired using the configuration listed in “**Optical setup for epi-fluorescent microscopy**” section. Images are presented by Photoshop and ImageJ. Images were analyzed by using CellProfiler as described in “Quantifying condensed fraction and condensate size, and colocalization by CellProfiler” section.

### Quantifying condensed fraction, condensate size, and colocalization by CellProfiler

To quantify *in vitro* condensates, two sets of images were taken under the same settings: One was buffer only, which is used for background fluorescence, and another was buffer plus proteins. The following pipeline was used to identify and measure condensates: identifying primary objects, measuring object intensity, measuring image intensity, and exporting to spread sheet. The parameters were used for identifying primary objects as follows: object size in pixel units 5 - 40; threshold strategy, global; threshold method, robust background; lower outlier fraction, 0.05; upper outlier fraction, 0.05; averaging method, mean; variance method, standard deviation; # of deviations, 20; threshold smoothing scale, 1.35; threshold correction factor, 1.0; and lower and upper bounds on threshold, 0 - 1.0. The condensed fraction was calculated as follows.

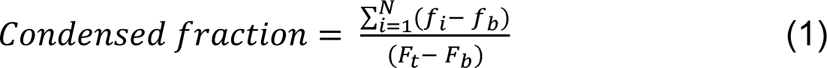

where *f_i_* is the fluorescence intensity of individual condensates, *f_b_* is the background fluorescence intensity of individual condensates, *F_i_* is the total fluorescence intensity of condensed and diluted parts, and *F_b_* is the total background fluorescence intensity.

The particle sizes of materials and aerosols have been well described by log-normal distribution (Alderliesten, 2016; Finlay and Darquenne, 2020; Heintzenberg, 1994; Patterson and Gillette, 1977; Waldo et al., 1971; Yang et al., 2012). The cumulative frequency distributions of condensate sizes in pixel were fitted with a lognormal cumulative distribution function based on the F-test implemented in OriginLab.

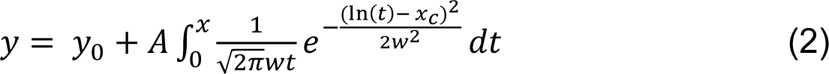

To quantify condensates of living cells, the following pipeline was used to identify and measure condensates: cropping a cell of interest; cropping a region without cells (background fluorescence); enhancing or suppressing features; identifying primary objects; measuring image intensity; measuring object intensity; and exporting to spreadsheet. The settings for identifying primary objects were as follows: object size in pixel units 2 - 20; threshold strategy, adaptive; threshold method, robust background; lower outlier fraction, 0.05; upper outlier fraction, 0.05; averaging method, mean; variance method, standard deviation; # of deviations, 1.01; threshold smoothing scale, 0; threshold correction factor, 1.02; lower and upper bounds on threshold, 0 - 1.0; and size of adaptive window, 10. The condensed fraction and condensate size were described as above. To quantify colocalization, the following pipeline was used: aligning, identifying primary objects of image 1; identifying primary objects of image 2; and measuring colocalization. The parameters for identifying primary objects were the same as described in “quantifying condensates of living cells”.

### Immunofluorescence

Approximately 0.5 × 10^6^ cells were seeded in each well of 6-well plate containing a cover glass and incubated overnight at 37°C, 5% CO_2_. The following day, the cells were washed with PBS and fixed with 1% formaldehyde (Sigma-Aldrich; P6148) for 10 minutes at room temperature. Cells were then washed with PBS and permeabilized by incubating with 0.2% Triton X-100 for 10 minutes with gentle shaking. The cells were incubated with blocking buffer (10.0 mM PBS pH 7.4, 0.1% Triton X-100, 3% Goat Serum, 3% BSA) for 2 hours. After washing with basic blocking buffer (10.0 mM PBS pH 7.4, 0.1% Triton X-100), the primary antibody was diluted in blocking buffer as follows: Anti-HaloTag (1:1000; Promega Corporation; G9281); anti-PHC1 antibody (1:500; Active Motif; 39723); anti-PHC2 antibody (1:500; Active Motif; 39661); Anti-RING1B (1:500; MBL; D139-3); anti-FLAG antibody (1:1000; Sigma; F1804), anti-BMI1 antibody (1:500; Cell Signaling; 6964S); and anti-H3K27me3 antibody (1:4000; Cell Signaling; 9733S). Cells were incubated with diluted primary antibodies for 2 hours at room temperature. After washing, secondary antibodies were diluted in blocking buffer as follows: Alexa Fluor 488-Goat anti-Mouse IgG (H+L) secondary antibody (1:1000; Life Technologies; A11029) an Alexa Fluor 568-Goat anti-Rabbit IgG (H+L) secondary antibody (1:1000; Life Technologies; A11011). After washing, cells were covered and incubated with diluted secondary antibodies for 2 hours at room temperature. After washing, the cover glass was mounted to a frosted microscope slide (Thermo Scientific; 2951-001) using ProLong^®^ Antifade reagents (Invitrogen; P7481) and left flat to dry overnight. Images were acquired using the configuration listed in “**Optical setup for epi-fluorescent microscopy**” section. Images were processed by Photoshop and ImageJ.

### FRAP

#### In vitro condensates

An 8-9 mm × 1.7 mm CoverWell™ Perfusion Chamber (Grace Bio-Labs; 622205) was adhered to a 25 × 25 mm micro cover glass. Perfusion chamber was filled with 100 µL of 0.1 mg/mL albumin and incubated for 10 minutes at room temperature. Albumin was then completely removed, and slide was immediately used for experiment. For the single-component system, 0.5 µM CBX2-YFP was incubated in the condensation formation buffer. For the two-component system, 0.5 µM of each of CBX2-YFP and RING1B-FLAG were mixed in the condensation formation buffer. For the three-component system, 0.5 µM of each of CBX2-YFP, RING1B-FLAG, and MEL18-FLAG were mixed in the condensation formation buffer. For the four-component system, 0.5 µM of each of CBX2-YFP, RING1B-FLAG, MEL18-FLAG, and PHC1-Cerulean were mixed in the condensation formation buffer. For the system containing Cy3-H3 K_c_27me3-nucleomses, the concentration of nucleosomes was 3.0 nM. After incubating for 10 min, samples were loaded into albumin-coated perfusion chamber for FRAP experiments as described below.

#### Live-cell condensates

mES cells with transgenic HT-CBX2, HT-CBX2 in *HT-CBX2/Ring1a^−/−^/Ring1b^fl/fl^* and *HT-CBX2/Mel18^−/−^/Bmi1^−/−^* were used for FRAP experiments. To deplete *Ring1b*, *HT-CBX2/Ring1a^−/−^/Ring1b^fl/fl^* cells were treated with 1.0 µM 4-hydroxytamoxifen (Sigma-Aldrich; H7904) for 48 hours prior to imaging. 3.0-5.0 × 10^5^ cells were seeded on a gelatin-coated 35-mm cover-glass bottom dish and incubated overnight at 37°C, 5% CO_2_. To visualize condensates, the mES cells were incubated with 50 nM HaloTag TMR ligand for 15 minutes. They were then washed with 2 mL PBS and incubated for another 30 minutes with 2 mL cell culture medium. This wash process was repeated a second time for a total of one hour of recovery. Finally, cells were loaded with imaging medium supplemented with 10% FBS and 10^3^ units/mL LIF for FRAP experiments as described below.

#### FRAP

Photobleaching was performed by using a Zeiss LSM 700 Observer with 100×/1.40 NA Oil-immersion Objective. The image size was 64.0 µm × 64.0 µm with an 8-bit image depth. For bleaching YFP, 488 nm laser was used. For bleaching Cy3 and TMR, 555 nm laser was used. The pinhole was fully open. Before photobleaching, two images were taken. After photobleaching, 30 images were taken with 5 second intervals. The images were analyzed by using ImageJ. TurboReg was used to correct for translational movement. The fluorescence intensities were corrected for fluctuations, and then normalized to the signal before bleaching as described previously (Ren *et al*., 2008; Zhen *et al*., 2014).

### Preparing cell and nuclear extract

For collection of whole-cell extract of HeLa cells stably expressing HT-PRC1 fusions with different Dox concentration treatments, cells (approximately 1 × 10^7^) were trypsinized and made single-cell by pipetting before being centrifuged at 300 × g for 5 minutes at 4°C. The cell pellet was resuspended in 200 µL of lysis buffer (20 mM Tris-HCl pH 7.4, 1.0 % IGEPAL^®^ CA-630, 200 mM NaCl, 0.25 mM EDTA, 20% glycerol, 0.1 mM Na_3_VO_4_, 0.5 mM DTT, 1× Protease Inhibitor Cocktail (Sigma; S8830), 0.1 mM PMSF) and rocked for 1.0 hour at 4 °C. Cell debris was pelleted by centrifugation and supernatant was collected. Since the expression of CBX2 is extremely low in mES cells, *Cbx2^HT/HT^* and *Cbx2^HT/WT^* knockin mES cells were cultured in the absence of LIF for two days and then nuclear extract was prepared. Approximately 2 × 10^8^ cells were incubated with hypotonic buffer (10 mM HEPES pH 7.5, 1.5 mM MgCl_2_, 10 mM KCl, and 0.1 mM PMSF) for 10 min and then struck slowly and steadily 12 times on ice by using type B pestle. Nuclei were collected by centrifugation and lysed with 200 µL of nuclear lysis buffer (20 mM Tris-HCl, pH 7.4, 2% NP-40, 300 mM NaCl, 0.25 mM EDTA, 20% Glycerol, 0.1 mM Na_3_VO_4_, 0.5 mM DTT, 1× Protease Inhibitor Cocktail, and 0.1 mM PMSF). After centrifugation, supernatant was collected.

### Western Blotting

Cell and nuclear extract were mixed with Laemmli Sample Buffer (Bio-Rad;1610747) and 10 mM DTT. After denaturing by heating to 95 °C for 10 min, samples were resolved in a NuPAGE™ 4-12% Bis-Tris Gel (ThermoFisher Scientific; NP0321BOX). The protein was transferred to an Immobilon^®^-FL PVDF Membrane (Millipore; IPFL00010) using a Trans-Blot^®^ SD Semi-Dry Transfer Cell (Bio-Rad; 1703940). After blocking. membrane was incubated with anti-HaloTag antibody (1:1000; Promega Corporation; G9281) and then with anti-Rabbit IgG Horseradish Peroxidase linked whole antibody (1:5000; GE Healthcare; NA943V). Membrane was visualized using Radiance Plus chemiluminescent HRP substrate (Azure Biosystems, AC2103) on Azure 300 imager (Azure Biosystems). After visualization, the membrane was stripped for 40 minutes with shaking using NewBlot™ PVDF Stripping Buffer (LI-COR; 928-40032). After blocking, membrane was incubated with anti-GAPDH (FL-335) rabbit polyclonal antibody (1:4000; Santa Cruz; SC-25778) and then with anti-Rabbit IgG Horseradish Peroxidase linked whole antibody (1:5000; GE Healthcare; NA943V). Membrane was visualized as described above.

### Coarse-grained simulations

The phase behaviour of proteins CBX2 and RING1B was investigated using a residue-level coarse-grained model for biomolecular phase separation that achieves quantitative agreement with experimental phase diagrams (Joseph *et al*., 2021). A modification was implemented to account phenomenologically for the presence of Mg^2+^ ions. In the model, each amino acid is represented via a single bead that has a unique charge, hydrophobicity, mass, and van der Waals radius. Within this framework, the energy of the system is computed as the sum of a scaled Wang-Frenkel potential for short-range pairwise interactions, coulombic Debye−Huckel term for long-range electrostatic interactions, and a standard harmonic potential for bonded interactions. The sequence of CBX2 and RING1B were the same as in the experiments. The initial atom-resolution structures for these proteins were obtained from AlphaFold. Globular domains were modelled as rigid bodies while disordered regions as fully flexible chains. To estimate the phase diagrams of CBX2, RING1B and the two-component mixtures, we use the Direct Coexistence method, where the protein-rich condensed phase and the protein-depleted dilute phase are simulated in the same simulation box separated by an explicit interface. For the single-component system, we used 54 copies of the protein (532 residues for CBX2 and 336 residues for RING1B), and for the two-component CBX2/RING1B mixtures, we used ZZ copies of each protein. Each system was first prepared in a cubic box. Isotropic *NPT*-ensemble (constant pressure and temperature) simulations were then performed at high pressure (>20 bars; using a Berendsen barostat) and low temperature (temperature regulated via a Langevin thermostat) to produce an initial high-density slab-like structure. One side of the box was then elongated (in the range of 3–10 times the box cross section) and *NVT*-ensemble simulations were then performed at several temperatures. Each system was simulated for 0.5 microseconds. We compute density profiles and monitor the energy of the systems to determine convergence. The presence of sharp interfaces is indicative of phase separation in our simulations, while the absence of an interface reveals that no phase separation occurs under a given set of conditions. Here, we report the temperature of our systems in terms of the critical temperature of the pure CBX2 systems. All simulations were performed using the LAMMPS simulation package. To estimate the critical point of the phase diagrams, we use the universal scaling law of coexistence densities near a critical point and the law of rectilinear diameters.

## ACKNOWLEDGMENTS

We acknowledge the Ren lab members for stimulating discussion during the writing process. This work was supported by the NIGMS under Award Number R01GM135286 (to X.R.) and the ORS at the University of Colorado Denver.

## AUTHOR CONTRIBUTIONS

X.R. and R.G. conceived and designed the study, supervised, and performed the experiments, analyzed data, prepared the figures, and wrote the paper. T.K. interpreted results and contributed to the writing of the manuscript. K.B, P.Y., S.I, J.E. and A.A. performed experiments, analyzed data, and wrote the manuscript.

